# Mechanism of homology search expansion during recombinational DNA break repair

**DOI:** 10.1101/2023.12.01.569403

**Authors:** Agnès Dumont, Nicolas Mendiboure, Jérôme Savocco, Loqmen Anani, Pierrick Moreau, Agnès Thierry, Laurent Modolo, Daniel Jost, Aurèle Piazza

## Abstract

Homology search catalyzed by a RecA/Rad51 nucleoprotein filament (NPF) is a central step of DNA double-strand break (DSB) repair by homologous recombination. How it operates in cells remains elusive. Here we developed a Hi-C-based methodology to map single-stranded DNA (ssDNA) contacts genome-wide in *S. cerevisiae*, which revealed two main homology search phases. Initial search conducted by short NPFs is confined in *cis* by cohesin-mediated chromatin loop folding. Progressive growth of stiff NPFs enables exploration of distant genomic sites. Long-range resection by Exo1 drives this transition from local to genome-wide search by providing ssDNA substrates for assembly of extensive NPFs. DSB end-tethering promotes coordinated homology search by NPFs formed on the two DSB ends. Finally, an autonomous genetic element on chromosome III engages the NPF and stimulates homology search in its vicinity. This work reveals the mechanism of the progressive and uneven expansion of homology search orchestrated by chromatin organizers, long-range resection, end-tethering, specialized genetic elements, and that exploits the stiff NPF structure conferred by Rad51 oligomerization.

**Highlights:**

– Cohesin-mediated chromatin loops constrain homology search in *cis* for NPF regions close to the resection front
– Stiffening of ssDNA by Rad51 enables genome-wide homology search by DSB-proximal sites
– Exo1-mediated long-range resection promotes genome-wide homology search
– DSB end-tethering promotes coordinated homology search by NPFs formed on both DSB ends
– The recombination enhancer focuses homology search in its vicinity

## Introduction

Templated DNA double-strand break (DSB) repair by homologous recombination (HR) entails a search for an intact homologous dsDNA templates in the excess of heterology of the genome and the nuclear volume. This search is conducted by a nucleoprotein filament (NPF) composed of RecA/Rad51 and additional proteins assembled along a hundreds-to kilobases-long ssDNA guide generated by the exonucleolytic resection of each DSB end ^1^.

The molecular mechanism of dsDNA sampling, mainly defined for bacterial RecA-ssDNA NPFs *in vitro* ^2^, occurs at the level of a few RecA monomers and is multiplexed along the length of a single NPF ^3,4^. The conservation of this inter-segmental search mode in eukaryote is supported by the ability for a single NPF to form D-loops and multi-invasion DNA joint molecules using internal regions of homology in *S. cerevisiae* cells, and with reconstituted yeast and human NPF *in vitro* ^5–9^. Accordingly, internal homologies are efficiently used as repair templates in *S. cerevisiae* ^10^. The whole length of the eukaryotic Rad51-ssDNA NPF is thus active in homology sampling and can lead to productive DNA strand invasion events, as observed for RecA *in vitro* ^11–13^.

A conserved property of RecA/Rad51-ssDNA NPFs is to adopt an elongated structure *in vitro*, with persistent length exceeding that of ssDNA by two to three orders of magnitude ^14–17^. Recent cytological observations with functional or recessive fluorescently-labeled RecA/Rad51 proteins supported the existence of similar, micrometer-long, stiff fiber-like structures in cells, which led to diverse models of homology search promotion in bacteria and eukaryotes ^18–21^. *E.coli* RecA-ssDNA NPFs stretching across the tubular nucleoid were proposed to reduce search complexity from 3D to 2D ^18^, while the search of comparatively shorter *C. crescentus* NPFs was promoted by cohesin-like RecN-mediated movements, presumably anchored within the RecA-ssDNA NPF ^21^. In *S. cerevisiae*, dynamic elongation and collapse of Rad51 structures were proposed to optimize exploration of a spherical nucleus ^20^. Here we sought to complement these cytological approaches by providing a population-averaged view of genomic sites engaged over time along unmodified Rad51-ssDNA NPFs on both sides of the DSB in *S. cerevisiae*.

Indeed, technical limitations in detecting intermediates of homology sampling by Rad51-ssDNA NPFs hampered progress in defining the homology search mechanism. Chromatin immunoprecipitation (ChIP) of Rad51 upon HO-induced DSB formation at the *MAT* locus on chr. III revealed homology-independent enrichment at the chromosome extremities, which was stimulated in a mating-type specific manner by the *cis*-acting recombination enhancer (RE) element ^22–24^. However, this method (i) precluded studying the role of factors affecting the formation and stability of the NPF, otherwise ablating its very readout. Furthermore, it could not resolve which segments of the Rad51-ssDNA NPF contacted other genomic sites. Differently, we recently leveraged Hi-C contact data of NPF-flanking dsDNA regions as a homology search readout ^25^, akin to DSB-flanking fluorescent operator-repressor arrays used to study the localization, association and mobility of DSB ends by cytological approaches ^26–30^. It revealed Rad51– and homology-dependent side-specific contacts between the DSB-flanking dsDNA and an inter-chromosomal donor site, confirming detection of *bona fide* recombinational interactions. However, this readout remains indirect and may not accurately reflect contacts made by NPF regions distant from the resection front.

We sought to overcome these limitations by developing a methodology derived from Hi-C that directly detects contacts made by ssDNA molecules. This ssDNA-specific Hi-C (ssHi-C) method revealed that homology search is not a uniform process along the NPF and the genome. It identifies various spatiotemporal features as well as genetic and protein determinants involved in regulating the gradual expansion of homology search from local to genome-wide.

## Results

### A Hi-C-based approach for detecting ssDNA contacts

Mapping of ssDNA contacts was undertaken using a well-characterized HO endonuclease-based system in *S. cerevisiae* ^31,32^. A single HO cut-site is located at *ura3*, 35 kb away left of the chr. V centromere and a 2 kb-long region of homology to the left DSB end is located at *lys2* on chr. II (**Figure 1A**). *HO* expression is induced upon S-release of G1-synchronized *MAT***a**-inc cells, yielding a highly homogeneous population of checkpoint-arrested cells attempting DSB repair by HR in G2/M at 2 and 4 hours post-DSB induction (**Figure 1B**). The efficient DSB formation in this system (>95% molecules cut in 1h) enables kinetics analysis of HR up to D-loop extension ^25,31,32^. To access the loci contacted by the DSB proximal regions, we performed Hi-C on cells dual-crosslinked with a short and a long-arm crosslinker (FA and EGS, respectively) upon chromatin digestion with DpnII and HinfI. This protocol yields high-quality stereotypic contacts maps at sub-kilobase resolution (median fragment length = 108 bp; see **Methods**) ^33^.

**Figure 1:**
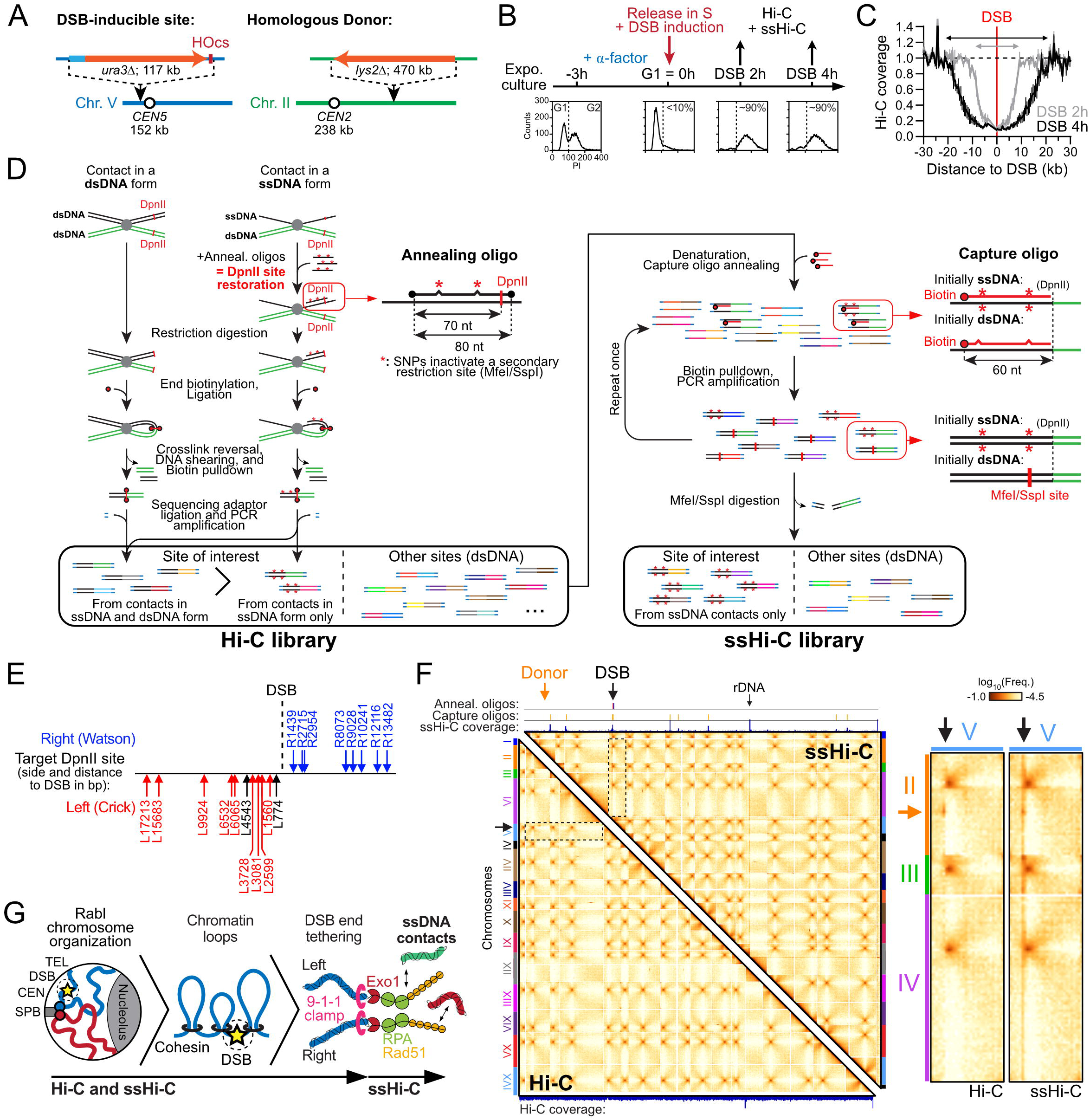
Genome-wide mapping of ssDNA and dsDNA contacts with ssHi-C. (A) Experimental system in haploid *S. cerevisiae*. (B) Cell synchronization, DSB induction and cell fixation protocol. >90% of cells have arrested in G2/M at 2 and 4 hours post-DSB induction. (C) Hi-C coverage following DSB induction, normalized on coverage in the absence of DSB. Data is mean±SEM of 4 biological replicates. (D) Rationale and steps of ssHi-C. (E) Annealing oligonucleotides used in this study. The black probes do not have corresponding capture oligonucleotides, and correspond to negative controls in (F). (F) Hi-C and ssHi-C contact maps of the whole genome 4 hours post-DSB induction in WT cells (APY266) showing classical features of yeast chromosome organization (*e.g.* centromere clustering and cohesin-mediated loop folding). Insets: Inter-chromosomal contacts between chr. V (black arrow) and chr. II to IV. A stripe of contacts emanating from the DSB site, with local enrichment visible at the donor site on chr. II (orange arrow) and on chr. III is only detected in the ssHi-C library. Absolute and relative coverage vs. undamaged cells show contact enrichment at dsDNA and ssDNA sites in ssHi-C. Maps are merge of 3 biological replicates subsampled at 19 million contacts. Bin: 20 kb. (G) ssHi-C retrieves yeast chromosome organization features in addition to ssDNA-specific contacts, allowing for the conjoined study of chromosome organization and homology search.

DSB resection produces long 3’-protruding ssDNA that eliminates restriction sites and the Hi-C readout along the region, resulting in a loss of signal (**Figure 1C**). To restore, isolate and enrich genome-wide contacts made by ssDNA, we devised a ssDNA-specific Hi-C procedure (hereafter ssHi-C) that relies on DpnII sites restoration upon annealing of 80mer oligonucleotides to the resected ssDNA (**Figure 1D**, **S1A** and **Methods**). These “annealing” oligonucleotides contain single-nucleotide polymorphisms (SNPs), some of which inactivate a secondary restriction site (MfeI or SspI), which will allow distinguishing contacts made at that site in the ssDNA or dsDNA form (**Figure 1D** and **S1A-B**). The Hi-C library is then subjected to two rounds of targeted enrichment for the ssDNA sites of interest using biotinylated “capture” oligonucleotides (**Figure 1D** and **S1A**). Digestion of the ssHi-C library with MfeI/SspI prior to sequencing eliminates contacts made at the sites of interest in a dsDNA form, thus isolating ssDNA contacts (**Figure 1D**). An additional SNP-based read filtering step further ensures isolation of ssDNA-specific contacts post-sequencing (**Figure S1B**, **Methods**).

We designed 21 annealing oligos targeting 19 DpnII restriction fragments on both sides of, and up to 17 kb away from, the DSB site. Out of the 19 ssDNA fragments, 17 are targeted by capture oligos, as well as 7 dsDNA sites used as positive enrichment controls and points of calibration between samples (**Figure 1E**). The two ssDNA fragments not captured are used as negative enrichment controls. ssDNA sites are labeled relative to the side (left or right) and the distance to the HO cut site (*e.g.* L2599 indicates a DpnII site at 2,599 bp to the left of the HOcs). One control dsDNA site (*ARG4*) is at the same distance from a centromere as the DSB site (**Figure S1C**). Annealing oligos partially restored Hi-C coverage, which could be specifically enriched using capture oligos (**Figure S2A-C**). No recovery and enrichment were observed at these sites in the absence of a DSB-inducible construct (**Figure S2A-D**), indicating that ssHi-C contacts are not produced by random ligation of annealing oligos, but instead require annealing to a cognate ssDNA. Site-to-site differences in ssDNA fragment enrichment partially depended on the fraction of resected DNA: the greater the resected fraction, the greater the ssDNA enrichment (**Figure S2D**). Some sites remained poorly restored, despite being fully resected (*e.g.* L1560 vs L2599, **Figure S2B**), suggesting site-specific differences in annealing efficiency. Various annealing oligo dropouts indicated that they (i) are each required, (ii) act independently, and (iii) do not interfere with one another for ssDNA site recovery (**Figure S2A, C**). In conclusion, ssHi-C recovers ssDNA-specific contacts for 16 out of the 17 fragments, with varying site-to-site efficiency.

Despite the strong enrichment for the 16 ssDNA and 7 dsDNA fragments targeted by the capture oligos, 87% of the total contacts originated from the non-targeted fragments. The resulting contact data were highly similar to that of conventional Hi-C (**Figure 1F** and **S2F-H**). The 7 control dsDNA sites recapitulated conventional Hi-C contacts, indicating that the capture procedure does not introduce bias at target and non-target sites (**Figure S2I-J**). ssHi-C libraries can thus be used to study both ssDNA contacts at the enriched sites as well as genome-wide dsDNA contacts classically detected with Hi-C (**Figure 1G**).

### The genome-wide distribution of ssDNA contacts is not uniform and departs from that of dsDNA in WT cells

Discrete, non-uniform stripes of contacts emanating from the DSB region stretched across chr. V and the genome (**Figure 1F** and **S2F**). No such stripes were observed at the 7 captured dsDNA regions, indicating that this contact behavior is specific to ssDNA sites and not an enrichment artefact (**Figure 1F**). A finer 4C-like representation reveals the propensity of ssDNA on both sides of the DSB to contact inter-chromosomal loci (hereafter *trans* contacts, 64% on average at 4 hours; **Figure 2A**). The frequency of ssDNA *trans* contacts increased significantly from 2 to 4 hours, an opposite trend to the chromatin condensation taking place concomitantly genome-wide ^25^ (**Figure 2B**). This *trans* contact frequency was significantly higher than that of (i) average dsDNA Hi-C *trans* contacts (19%), (ii) DSB-flanking dsDNA Hi-C *trans* contacts (33.7%), and (iii) a control captured dsDNA site located at the same distance from a centromere as the DSB site (*ARG4*, 22%; **Figure 2A-B** and **S3A-B**). The distribution of ssDNA-specific *trans* contacts was not uniform over the genome, with local enrichment at centromeres, the donor site, and on chr. III (**Figure 2A** and see below). Finally, the distribution of *trans* contacts was not uniform along ssDNA, from ∼74% for DSB-proximal sites to ∼40% for DSB-distal sites 4 hours post-DSB induction (**Figure 2C**). Consequently, the predominant ssDNA contacts with chromatin in *trans* is a decreasing function of the genomic distance to the DSB in wild-type cells. These observations show that chromatin engagement by ssDNA is not uniform over the genome, along ssDNA, and over time.

**Figure 2:**
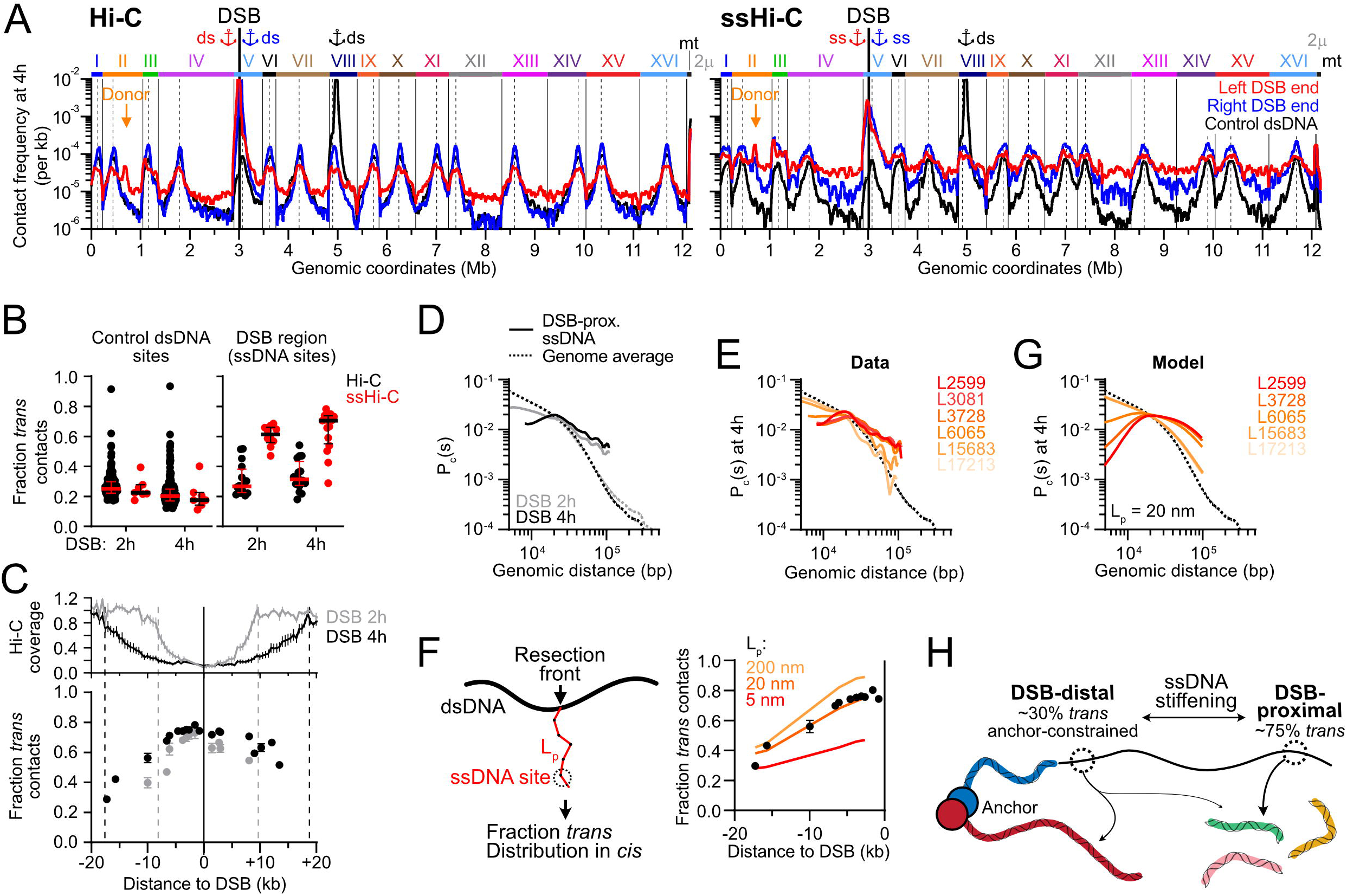
DSB-proximal ssDNA sites emancipate from *cis* spatial constraints over time. (A) Average genome-wide contact distribution of the left and right DSB ends determined with Hi-C (left; dsDNA only, viewpoint 5 kb away from the DSB site) and ssHi-C (right; ssDNA only, average of all ssDNA sites) 4h post-DSB induction in WT cells (APY266). A control dsDNA site on chr. VIII, located at the same distance to a centromere as the DSB (**Figure S1C**), is shown for comparison. Dotted lines: centromeres. Anchors: viewpoints. Orange arrow: donor. Data represent the average of 4 biological replicates smoothed over 30 kb. (B) *Trans* contacts for dsDNA and ssDNA sites determined in the DSB region (right) and at non-damaged sites genome-wide (left) with Hi-C and ssHi-C at 2 and 4 hours post-DSB induction in WT cells. Data points: individual ssDNA sites (ssHi-C) or 1 kb dsDNA bins (Hi-C). (C) Top: Hi-C coverage. Bottom: Proportion of ssDNA contacts in *trans* as a function of the distance to the DSB. Dotted lines denote the position of the resection fronts. (**E**) LOWESS regression of average contacts made by ssDNA sites on the left DSB end in *cis* compared to the genome-averaged P_c_(s) at 2 and 4 hours post-DSB induction. Dotted line: Genome-average dsDNA P_c_(s). (**G**) LOWESS regression of individual contacts made by ssDNA sites on the left DSB end in *cis* at 4 hours post-DSB induction compared to the genome-averaged dsDNA P_c_(s) (dotted line). Data are from (A). (**H**) Left: “Fishing line” modeling rationale. The red polymer represents ssDNA emanating from the dsDNA polymer. L_p_: persistence length. Right: Predicted fraction of *trans* contacts at increasing distance from the resection front with different L_p_. (**I**) Predicted distribution of *cis* contacts for sites in (G) with L_p_ = 20 nm. (**J**) ssDNA stiffening projects DSB-proximal ssDNA sites away from DSB-distal sites and the initial chromatin environment.

### DSB-proximal ssDNA contacts with chromatin in *cis* are broadly distributed

The distribution of *cis* contacts made by ssDNA sites departed from that of flanking dsDNA regions detected with Hi-C (**Figure 2A** and **S3C**). First, each ssDNA end similarly contacted chromatin on both sides of the DSB, while contacts by DSB-flanking dsDNA regions were asymmetrically distributed across the DSB region (**Figure S3C**). Second, ssDNA contacts enrichment with chromatin in *cis* decreased from 2 to 4 hours, coincident with an increase in *trans* contact (**Figure 2C** and **S3C**). Third, the decay of ssDNA contacts with genomic distance did not follow the genome-average probability of dsDNA contacts as a function of genomic distance (P_c_(s)), but instead appeared to decay more slowly (**Figures 2D**). Examination of individual ssDNA sites revealed that DSB-proximal ssDNA contacts were broadly distributed in *cis*, enriched 10-fold at the highest over the genome average (**Figure S3D**). In contrast, DSB-distal ssDNA sites exhibited a strong contact enrichment with DSB-flanking chromatin, which decayed following the genome-averaged P_c_(*s*) (**Figure 2E** and **S3D**). More specifically, the proximity to the resection front correlated with frequent ssDNA contacts with chromatin in *cis*, as well as across the resected region (*e.g.* the R8073 site, at the resection edge at 2 hours and ∼8 kb away at 4 hours post-DSB induction, **Figure S3E**). These observations suggest that ssDNA sites are progressively relieved from constraints confining most contacts in *cis* over time, which correlates with the increasing distance to the resection front (see below).

### DSB-proximal and DSB-distal ssDNA regions are differently emancipated from a *cis*-acting spatial anchor

The Rabl organization of yeast chromosomes is defined by the anchoring of centromeres at the spindle-pole body (SPB) via microtubules and of telomeres at the nuclear envelope, whose distance from the SPB scales with chromosome arm length ^34–39^. This organization spatially constrains surrounding chromatin motion, promotes clustering between ∼100 kb centromeric regions and inhibits centromere-telomere contacts ^37,40–44^ (**Figure 1F**). The DSB site, positioned 35 kb left of *CEN5*, is subjected to such constraint (**Figure 1A**). Accordingly, ssDNA sites preferentially engaged centromeres in *trans*, especially for *CEN5*-associated DSB-distal ssDNA sites, yet to a lower extent than undamaged dsDNA sites (**Figure 2A** and **S3F**). Conversely, DSB-proximal ssDNA sites engaged telomeres of long chromosome arms more frequently than DSB-distal sites (**Figure S3G**). These contact profiles depended on centromere clustering at the SPB, as treatment with the microtubule-depolymerizing drug nocodazole (i) reduced ssDNA contacts with centromeres, (ii) largely relieved the inhibition of interactions with telomeres of long chromosome arms, and (iii) abolished the distinction between DSB-proximal and –distal ssDNA sites, especially for *CEN5*-associated ssDNA sites (**Figures S3H-M**). These results show that DSB-proximal ssDNA is less constrained by a spatial anchor relative to a DSB-distal ssDNA sites.

### ssDNA stiffening projects DSB-proximal sites away from the resection front and the initial chromatin environment

To interpret the observed changes in *cis* and *trans* contacts at ssDNA sites at varying distances from the DSB, we propose a theoretical framework where we modeled the NPF as a semi-flexible polymer – characterized by a persistence length L_p_ – emanating from the resection front and fishing for contacts (**Figure 2F**, see **Methods** and **Supplementary Code 1** for details). Using a simple Gaussian contact probability model coupled to a stochastic description of resection, we computed the expected fraction of *trans* contacts and the distribution of *cis* contacts as a function of the distance between the ssDNA site and the DSB (**Figure 2F, Supplementary Code 1**). More rigid NPFs leads to an increase in the *trans* fraction and to a relative increase in long-range *cis* contacts compared to local *cis* ones. Given the typical contour length of an ATP-bound RecA/Rad51 filament (∼0.5 nm per nt of ssDNA)^15^ and the inferred dynamics of the resection front (speed ∼3.9 kb/h), the value L_p_ ≈ 20 nm yielded *trans* contacts and *cis* contact profiles in good quantitative agreement with that observed 4 hours post-DSB induction (**Figure 2G**). This apparent L_p_ is at least an order of magnitude greater than that of protein-free or RPA-coated ssDNA *in vitro* ^45–48^. It indicates that ssDNA is stiffened in the context of DSB repair by HR in cells, leading to the distinct contact profiles of DSB-proximal and DSB-distal ssDNA sites (**Figure 2H**).

### ssDNA stiffening by Rad51 promotes genome-wide homology search

We evaluated the contribution of Rad51 in regulating ssDNA stiffness and genome-wide ssDNA contacts at 2 and 4 hours post-DSB induction in a *rad51Δ* mutant. Absence of Rad51 did not markedly affect resection, DSB end-tethering, metaphase-arrest, cohesin-mediated loop folding, chromosome individualization, or the Rabl chromosome organization ^25^ (**Figure 3A** and **S4A-E**).

**Figure 3:**
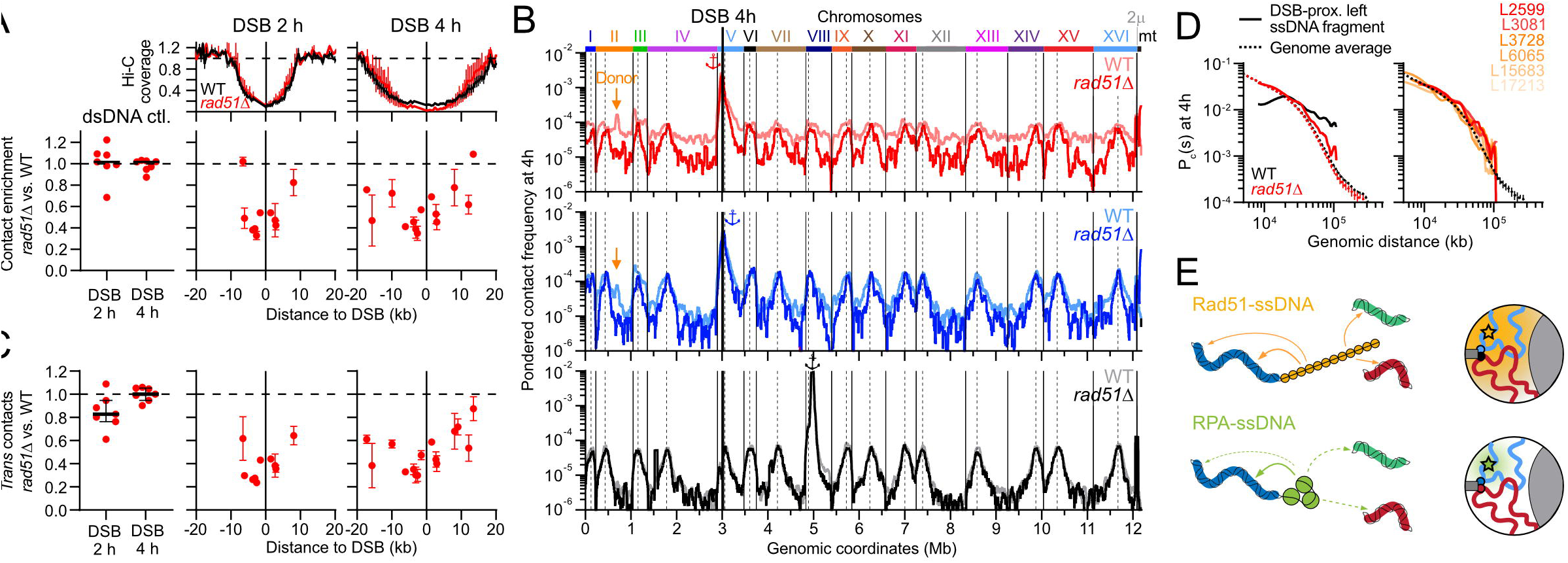
Rad51 promotes genome-wide homology search by DSB-proximal ssDNA sites by stiffening ssDNA. (A) Hi-C coverage (top) and amount of dsDNA (left) and ssDNA (right) contacts retrieved in a *rad51Δ* mutant (APY679) relative to WT cells (bottom). Data: mean ± SEM of individual captured ssDNA and dsDNA sites from 2 biological replicates. (B) Average genome-wide contact distribution of the left and right ssDNA sites 4 hours post-DSB induction in WT cells (APY266) and *rad51Δ* cells (APY679). Other legends as in Figure 2A. (C) *Trans* contact frequency relative to WT cells for individual ssDNA sites (right) and 7 control dsDNA sites (left) at 2 and 4 hours post-DSB induction in *rad51Δ* mutants in (B). Error bars show SEM. (E) LOWESS regression of the average (left) and individual (right) *cis* contacts made by ssDNA sites on the left DSB end, compared to the genome-averaged P_c_(s) in WT and *rad51Δ* cells. (F) Genome-wide homology search by DSB-proximal ssDNA sites is promoted by stiff Rad51-ssDNA filaments.

The two DNA binding sites of RecA/Rad51 bridge the ssDNA guide and a dsDNA molecule within synaptic complexes, unlike RPA. Once crosslinked, such Rad51-mediated ssDNA-dsDNA association will lead to the ligation events reported by Hi-C ^49^. Consistently, the amount of captured ssDNA fragments decreased 2-to 3-fold in a *rad51Δ* mutant compared to WT cells (**Figure 3A**). Calibration of the contact distribution on the capture efficiency of each ssDNA fragment enabled determining the absolute gain and loss of contacts relative to the WT context (**Methods**). Strikingly, *trans* contacts decreased ∼3-fold in the absence of Rad51 for DSB-proximal ssDNA sites, back to levels observed for dsDNA sites (**Figure 3B, C** and **S4F**). Long-range *cis* contacts decreased while short-range (<20 kb) contacts increased modestly or remained constant on both sides and across the DSB region (**Figure S4G**). Contacts in *trans* decreased everywhere except at the spatially-associated centromeres (**Figure 3B** and **S4F, I-J**). Even when considering only *trans* contacts, interactions with long-arm telomeres decreased to levels observed for intact dsDNA sites (**Figure S4J**). Contacts with the donor site were abolished (**Figure 3B** and **S4F**). Consequently, the engagement of distant chromosomal regions by ssDNA ends are not due to the rupture of the dsDNA polymer, but instead reflects ongoing homology search by the Rad51-ssDNA NPF.

Unlike in WT cells, the fraction of *trans* contacts as well as the distribution of contacts in *cis* remained similar for all ssDNA sites (**Figure S4H**) and followed the genome-averaged P_c_(s), irrespective of their distance to the DSB end in *rad51Δ* cells (**Figure 3D**). Distant ssDNA sites are thus spatially associated and at a null distance from the resection front in the absence of Rad51. These observations show that Rad51 extends the ssDNA molecule in WT cells, which otherwise collapses upon RPA binding in *rad51Δ* cells (**Figure 3E**), an interpretation consistent with globular RPA-ssDNA structures and elongated Rad51/RecA-ssDNA structures observed *in vitro* and in cells ^14,16,18–21,48,50–54^. Consequently, stiffening of ssDNA by Rad51 projects DSB-proximal sites away from the DSB-borne chromatin environment, which promotes genome-wide homology search (**Figure 3E**).

### Exo1-dependent generation of long ssDNA promotes genome-wide homology search

Since long-range resection produces extensive ssDNA for NPF assembly, we addressed its contribution to genome-wide homology search by determining the distribution of ssDNA contacts in *exo1* mutants^1^. The *exo1Δ* and a catalytic-deficient *exo1-D173A* (hereafter *exo1-cd*) mutant gave highly similar results and were pooled (**Figure S5A-G** and see below). The rate of long-range resection was reduced from ∼3.9 kb/h in WT cells to ∼1.5 kb/h in Exo1-deficient cells (**Figure 4B**, **Supplementary Code 1**). As previously noted, Exo1 deficiency also caused a loss of DSB end-tethering, which resulted in the detachment of the centromere-devoid chromosomal fragment ^25,55^ (**Figure 4A and S5A-G**). However, the Rabl organization, metaphase-arrest, cohesin-mediated loop organization and chromosome individualization were not affected in Exo1-deficient cells ^25^ (**Figure 4A** and **S5A-E**).

**Figure 4:**
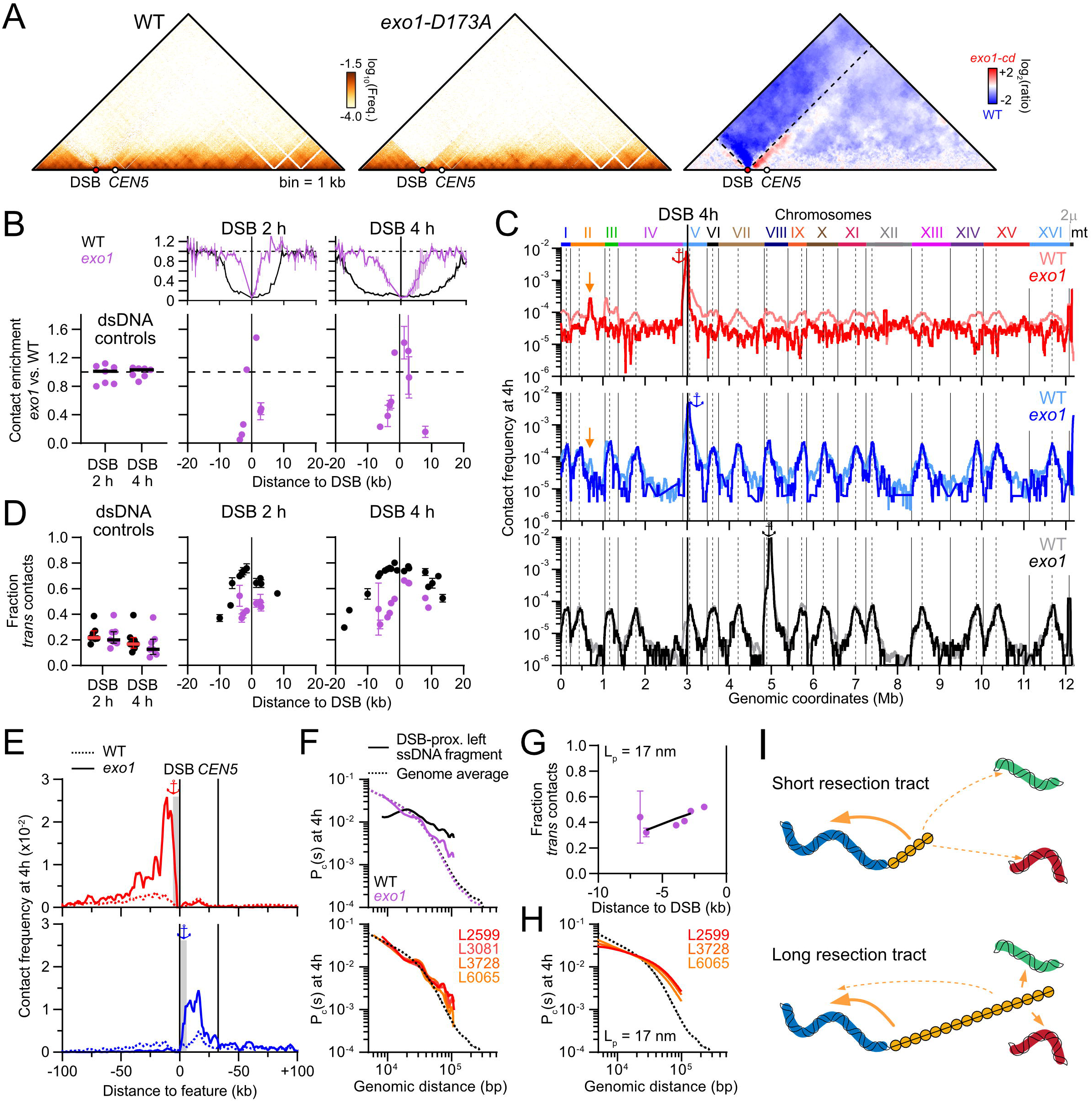
Exo1 catalytic activity promotes genome-wide homology search. (A) Left: Hi-C contact maps of chr. V in WT (APY266) and *exo1-D173A* (APY536) cells 4 hours post-DSB induction. Right: Ratio map of the *exo1-D173A* over WT contacts using Serpentine binning ^94^. (B) Hi-C coverage (top) and amount of dsDNA (left) and ssDNA (right) contacts retrieved in *exo1Δ* and *exo1-D173A* mutants (APY1160 and APY536, respectively) relative to WT cells (bottom). Data points: mean ± SEM of individual captured ssDNA and dsDNA sites from a pooling of both mutants (one biological replicate each). (C) Average genome-wide contact distribution of the DSB-proximal left and right DSB ends 4h post-DSB induction in WT cells (APY266) and pooled Exo1-deficient cells (APY1160 and APY536). Other legends as in Figure 2A. (D) Average *trans* contact frequency for individual ssDNA sites (right) and 7 control dsDNA sites (left) at 2 and 4 hours post-DSB induction in WT (n=4) and Exo1-deficient (n=2) cells in (C). Error bars show SEM. (E) Zoom of the ssHi-C data in (C) around the DSB region, smoothed over 4 kb. (F) LOWESS regression of the average (top) and individual (bottom) *cis* contacts made by ssDNA sites on the left DSB end, compared to the genome-averaged P_c_(s) in WT and Exo1-deficient cells. (**G-H**) Predicted fraction of *trans* contacts (G) and distribution of *cis* contacts (H) at increasing distance from the DSB site with L_p_ = 17 nm compared to experimental data from (D). (**I**) Model for the Exo1-driven genome-wide expansion of homology search.

As expected, Exo1 deficiency led to a loss of ssDNA contacts at DSB-distal regions as they remained non-resected (**Figure 4B**). In contrast, resected DSB-proximal ssDNA sites yielded contacts enrichment equivalent to WT (**Figure 4B**), indicating no notable defect in dsDNA engagement by the NPF in *exo1* mutants. The distribution of ssDNA contacts was strongly altered in Exo1-deficient cells. First, the left NPF contacted the genome in a uniform manner, owing to the loss of end-tethering in these mutants (**Figure 4C** and **S5G-H**). Contacts between the right NPF and centromeres were exacerbated in Exo1-deficient mutants compared to WT cells (**Figure 4C** and **S5H**). Second, NPF were defective in engaging dsDNA in *trans* (**Figure 4D**). Third, among this reduced fraction of *trans* contacts, interaction with long-arm telomeres by the centromere-bound NPF were depleted, similar to undamaged dsDNA sites (**Figure S5I**). These observations indicate a defect for the NPF in contacting distant dsDNA sites.

Conversely to this decrease of *trans* contacts, ssDNA contacts by both DSB ends increased with chromatin in *cis* in a side-specific manner (**Figure 4E** and **S5J**). The *cis* contact profiles followed the genome-averaged P_c_(s) more closely than in WT cells (**Figure 4F**). They also differed from that observed in a *rad51Δ* mutant, in which all sites exhibited similar slopes to the genome-averaged P_c_(s) (**Figure 3E**). The Gaussian contact probability model satisfyingly accounted for both the fraction of *trans* contacts and the distribution of *cis* contacts with L_p_ ≈ 17 nm (**Figure 4G-H**), very similar to the persistence length obtained in WT cells (L_p_ ≈ 20 nm, **Figure 2H-I)**. Consequently, NPF properties are not altered in Exo1-deficient cells.

Finally, the slope of *trans* contacts as a function of the distance to the right resection front was similar for Exo1-deficient and WT cells (**Figure S5K;** detachment of the left chromosome fragment prevented a straightforward comparison of the contacts made by the left DSB end). The increased propensity to access genomic regions in *trans* thus reflects the genomic distance between a ssDNA site and the resection front, not the time spent as ssDNA. Altogether, these observations show that long-range resection mediated by Exo1 progressively drives genome-wide homology search by providing increasingly long ssDNA substrate for assembly of Rad51-ssDNA NPF (**Figure 4I**).

### Cohesin-mediated chromatin looping focuses search by nascent Rad51-ssDNA NPFs within flanking chromatin loops

We previously showed that cohesin-mediated loop folding inhibits homology search in *trans* using DSB-flanking dsDNA Hi-C contacts and quantification of D-loop formation of a DSB-proximal sequence at *cis* and *trans* ectopic donors ^25^. This inhibition was more pronounced at 2 hours than at 4 hours post-DSB induction, which suggested a decreasing influence of cohesin-mediated loop folding on homology search by DSB-proximal sites over time ^25^. We addressed the role of cohesin in restricting genome-wide ssDNA contacts upon auxin-induced depletion of the Scc1 subunit of cohesin ^25^ (**Figure 5A**). As shown previously, Scc1 depletion abolishes sister chromatid cohesion, chromatin loop folding and chromosome compaction while retaining the checkpoint-mediated G2/M-arrest ^25,56,57^ (**Figure S6A-D**). Partial Scc1 depletion relieved an inhibition on *trans* contacts by all ssDNA sites, without markedly affecting the overall ability of the NPF to engage dsDNA (**Figure 5B** and **S6E-G**). No such relief was observed in a *rad51Δ* mutant background, which again exhibited a defect in contacting dsDNA (**Figure 5B** and **S6G**). Consequently, cohesin inhibits NPF interactions in *trans.* This cohesin-mediated inhibition was particularly pronounced for DSB-distal NPF sites, while DSB-proximal sites exhibited comparable levels of *trans* contacts at both 2 and 4 hours post-DSB induction (**Figure 5B**). Scc1 depletion did not affect the distribution of *cis* contacts by DSB-proximal sites, which extended at much longer range than the genome-wide P_c_(s) in a Rad51-dependent manner, as in WT cells (**Figure 5C** and **S6H**). Similarly, DSB-distal sites behaved as in WT cells and followed the genome-averaged P_c_(s) upon Scc1 depletion (**Figure 5C** and **S6H**). Consequently, cohesin inhibits long-range *cis* and *trans* homology search by DSB-distal sites within ∼10 kb from the resection front.

**Figure 5:**
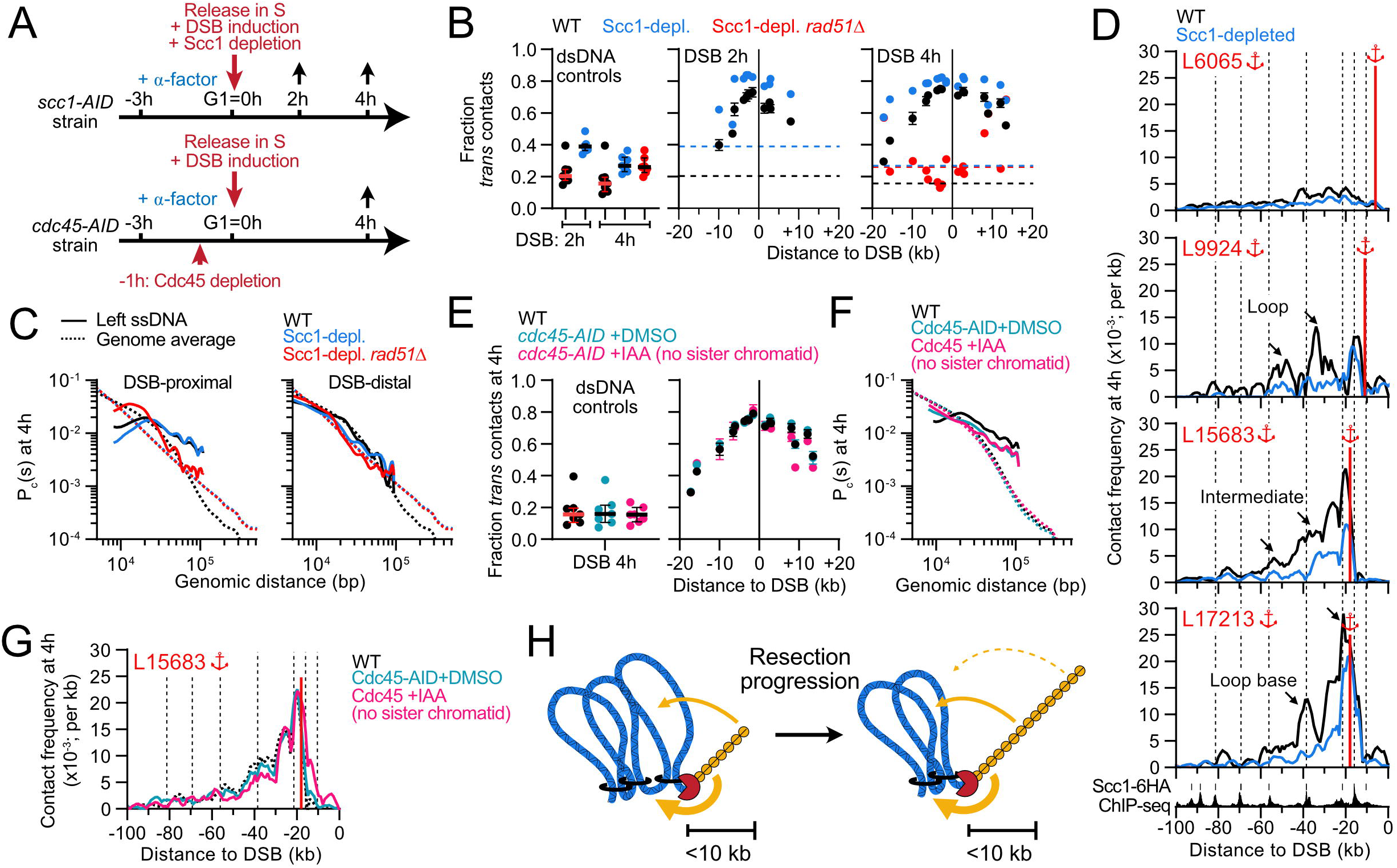
Cohesin directs homology search by DSB-distal ssDNA sites in *cis*. (A) Experimental schemes for depletion of AID-tagged Scc1 and Cdc45. (B) Average *trans* contact frequency for individual ssDNA sites (right) and 7 control dsDNA sites (left) at 2 and 4 hours post-DSB induction in WT (n=4), Scc1-depleted cells (APY1481) and Scc1-depleted *rad51Δ* cells (APY1500). (C) LOWESS regression of the average (top) and individual (bottom) *cis* contacts made by DSB-proximal (left panel) and DSB-distal (right panel) ssDNA sites on the left DSB end, compared to the genome-averaged P_c_(s) in WT, Scc1-depleted and Scc1-depleted *rad51Δ* cells. (D) Side-specific *cis* contact profiles of individual ssDNA sites in WT and Scc1-depleted cells 4 hours post-DSB induction. The calibrated Scc1-6HA ChIP-Seq profile obtained in this system 4h post-DSB induction in WT cells are from ref. ^25^. Dotted lines: position of cohesin peaks. Arrows: cohesin-dependent enrichment of ssDNA contacts. (E) Average *trans* contact frequency for individual ssDNA sites (right) and 7 control dsDNA sites (left) at 4 hours post-DSB induction in untreated WT cells (n=4) and Cdc45-AID-tagged cells (APY513) either mock-treated or treated with IAA (n = 2 each) as depicted in (A). (F) LOWESS regression of the average *cis* contacts made by DSB-proximal ssDNA sites on the left DSB end compared to the genome-averaged P_c_(s) in WT, Cdc54-tagged and Cdc45-depleted cells. (G) Side-specific *cis* contact profiles for the DSB-distal L15683 ssDNA site in WT, Cdc54-tagged and Cdc45-depleted cells 4 hours post-DSB induction. Other legends as in (D). (H) Model for the interaction of DSB-distal ssDNA sites with side-specific cohesin-mediated chromatin loop in *cis*. Newly generated ssDNA interacts with loop base, which are also contact points between sister chromatids ^72^. This configuration presumably puts nascent NPF in close proximity to the sister chromatid. Resection progression projects these sites at increasing distance from loop base over time.

Examination of individual ssDNA sites further revealed that the NPF region closest to the resection front contacted nearby chromatin predominantly at cohesin-enriched sites (L9924 and L17213 at 2 and 4 hours post-DSB induction, respectively; **Figure 5D** and **S6I**). NPF sites further away from the resection front progressively contacted the chromatin located between cohesin peaks, *i.e.* the inner part of chromatin loops (L6065 at 2 hours and L15683 and L9924 at 4 hours; **Figure 5D** and **S6I**). This observation was also true for DSB-distal sites on the right DSB end, although limited to a single loop flanking the centromere of chr. V (**Figure S6J**), possibly owing to the inability for cohesin to cross the centromere ^57,58^. Finally, DSB-proximal sites exhibited little cohesin-dependent *cis* contact enrichment (**Figure 5D** and **S6K**). Cohesin also promoted interactions across the DSB region (**Figure S6I**). The contact profile appeared broad without clear relation with the position of cohesin-enriched sites, even for ssDNA sites close to the resection front, unlike side-specific contacts (**Figure S6J**). These observations show that cohesin directs homology search in *cis* by DSB-distal ssDNA sites in a side-specific manner.

Whether the role of cohesin involved sister chromatid cohesion was addressed upon depletion of the replication factor Cdc45 prior to releasing cells from their G1 arrest ^59^ (**Figure 5A**). Cdc45-depleted cells reach G2/M with unreplicated chromosomes but retain chromatin loop folding similar to that observed in WT cells ^25,57^. The same cell pellets as those used and controlled for in ref. ^25^ were re-processed for ssHi-C. Cdc45 depletion did not relieve the cohesin-mediated inhibition of NPF *trans* contacts, nor did it affect contacts with chromatin in *cis* **(Figure 5E-G**). Consequently, the mechanism by which cohesin regulates DSB-distal NPF contacts is independent of its role in sister chromatid cohesion.

These observations extend our prior conclusions that cohesin-mediated loop folding both (i) inhibits homology search in *trans* upon chromosome individualization, and (ii) promotes homology sampling in chromatin loops extruded near the resection front in *cis* ^25^ (**Figure 5H**). Moreover, our new results show that this influence is exerted primarily over a ∼10 kb-long NPF regions flanking the resection front. Consequently, DSB-proximal sites overcome the cohesin-mediated constraint to genome-wide homology search upon long-range resection progression and NPF growth.

### The Recombination Enhancer focuses homology search on chr. III

Intriguingly, chr. III was disproportionately engaged by ssDNA sites on both sides of, and at all distances from the DSB (**Figure S7A**). This enrichment was most notable 2 hours post-DSB induction, indicative of an early HR event (**Figure S7A**). Other small chromosomes (I and VI) were significantly less contacted than chr. III and contact enrichment with chr. III was retained upon nocodazole treatment, ruling out the outsize influence of centromeres clustering on short chromosomes as the sole determinant for this enrichment (**Figure S7B**). Contacts by both ssDNA ends were mainly focused in a ∼30 kb region surrounding the Recombination Enhancer (RE), enriched by at least an order of magnitude above average inter-chromosomal contacts (**Figure 6A**). The RE is a discrete genetic element that stimulates HR repair with donor sequences located in its vicinity, inside and outside of the context of mating-type switching ^60–62^. Contact enrichment was also notable in a broader ∼100 kb region on the right end side of chr. III (**Figure 6A**), presumably owing to the RE-dependent horseshoe folding of chr. III ^42,63,64^ (and see below **Figure 6C**). Rad51 is required for this ssDNA-RE interaction, indicating that these contacts involve the NPF (**Figure 6B** and **S7C**). The NPF-RE interaction did not require cohesin (**Figure S7D**). However, it depended on the catalytic activity of Exo1 (**Figure S7E, F**), but whether it pertains to its role in forming extended NPFs, in end-tethering, or in regulating the DNA damage checkpoint ^65,66^ is unclear (see **Discussion** below). Deletion of the RE abolished contact enrichment on chr. III (**Figure 6C**). Conversely, addition of the minimal 700 bp-long RE ^67^ in the middle of the 1 Mb-long right arm of chr. IV promoted contacts between the NPF and the ∼30 kb surrounding region (**Figure 6D** and **S7G**). Consequently, the RE is required and sufficient for NPF recruitment.

**Figure 6:**
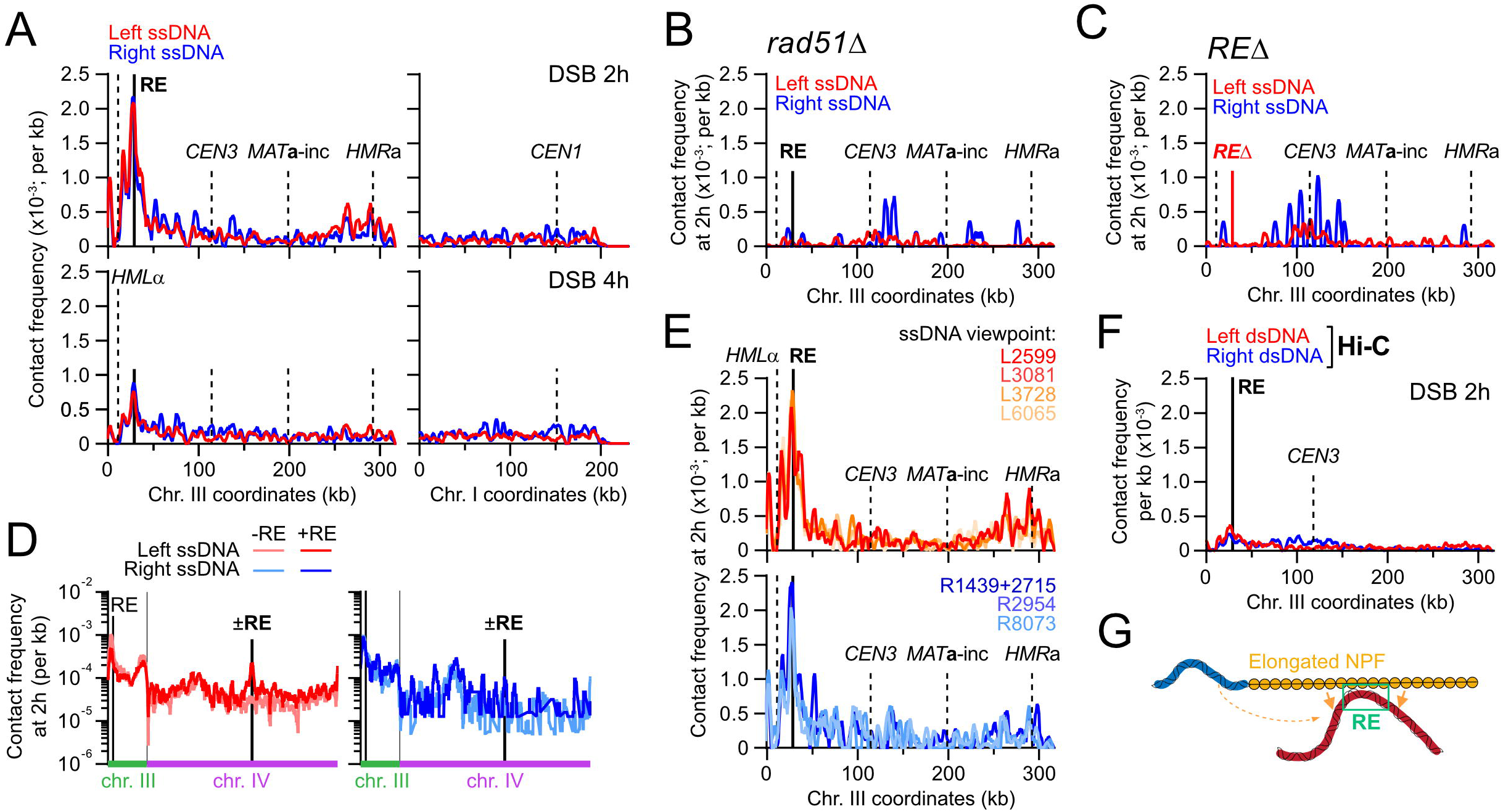
The recombination enhancer stimulates homology search in *trans* on chr. III. (A) ssHi-C contact profiles on chr. I and III at 2 and 4 hours post-DSB induction in WT cells. From data in Figure 2. (B) Pondered ssHi-C contact profiles on chr. III in *rad51Δ* cells 2 hours post-DSB induction. (C) ssHi-C contact profiles on chr. III in a strain deleted for the RE (APY1548) 2 hours post-DSB induction. (D) ssHi-C contact profiles on chr. III and IV in WT cells (APY266) and in cells bearing an additional 700 bp RE element inserted at position 845,464 on chr. IV (APY1497) 2 hours post-DSB induction. Signal smoothed over 10 kb. (E) ssHi-C contact profiles of individual ssDNA sites on the left and right DSB ends. Signal smoothed over 4 kb. (F) Hi-C contact profiles of DSB-flanking dsDNA regions 5 kb away to the left and right of the DSB site in WT cells 2 hours post-DSB induction. (G) Model for the NPF-RE interaction.

DSB-proximal and DSB-distal NPF segments equally contacted the RE (**Figure 6E**). In contrast, DSB-flanking dsDNA regions contacted the RE ∼10-fold less than NPFs did (**Figure 6F**). These observations suggest that (i) all regions along the Rad51-ssDNA NPF are equally prone to bind to the RE, and (ii) that this NPF-RE binding takes place in an extended NPF, on average at a distance from the flanking dsDNA chromatin (**Figure 6G**), unlike NPF-donor interactions (see below).

These observations reveal the existence of a mechanism for local homology search enhancement upon direct interaction between a specialized genetic element and the Rad51-ssDNA NPF that can operate in *trans*.

### Homology search is partly coordinated between NPFs on both DSB ends

The contact profiles of the left and right DSB ends are not super-imposable, yet they differ less than upon loss of DSB end-tethering in Exo1-deficient cells, suggesting that Rad51-ssDNA segments are associated a substantial fraction of the time, or in a substantial fraction of the cell population, during homology search ^20^ (**Figure 2A, 4C,** and **4E**). In order to address this possibility, we focused on contacts with the donor site, which has homology with the left DSB end only. Both left and right NPFs interacted with a ∼60 kb region surrounding the donor site, in a homology– and Rad51-dependent manner (**Figures 7A-C**, and **S8A**). The homology-containing NPF to the left of the DSB contacted the donor region on average 2-to 3-fold more frequently than the heterologous right NPF (**Figure 7A**). DSB-proximal NPF regions exhibited even smaller differences, only ∼1.5-fold higher for the left than the right NPF (**Figure 7B**), suggesting that NPF are associated in a co-oriented (*i.e.* 5’-3’) configuration. Loss of DSB end-tethering in Exo1-deficient cells caused a loss of contacts specifically between the heterologous right NPF and the donor (**Figure 4C, 7C**, and **S8A**). Consequently, left-right NPF interactions are not reestablished once homology has been found by one end. It instead indicates that the majority of the NPFs are associated and carry out homology search in a coordinated manner.

**Figure 7:**
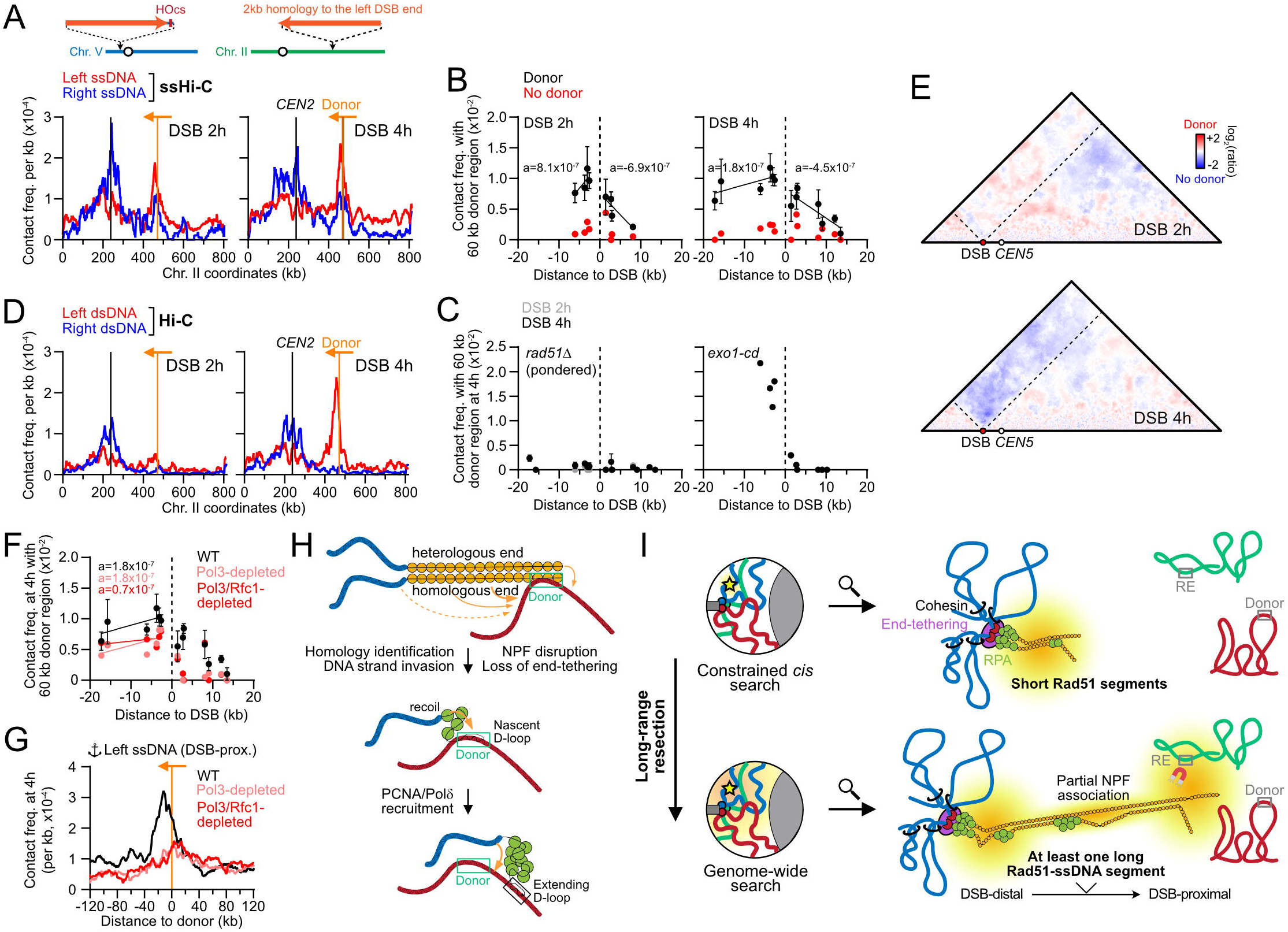
Coordinated homology search by both DSB ends and side-specific NPF collapse in a donor-dependent manner. (A) ssHi-C contact profiles on chr. II in WT cells (APY266). From data in Figure 2. Orange bar and arrow: position and orientation of the homologous region to the left DSB end. Data smoothed over 10 kb. (B) Contact frequency of individual ssDNA sites with a 60 kb region surrounding the donor site on chr. II in the presence (APY266; n = 4) and absence (APY358, n = 1) of homology for the left DSB end at that site. a: slope of the linear regression on both side of the DSB. Data: mean ± SEM. (C) Same as (B) in a *rad51Δ* mutant (left) and an *exo1-D173A* mutant (right). (D) Hi-C contact profiles on chr. II of DSB-flanking dsDNA regions 5 kb away to the left and right end of the DSB site. Other legends as in (A). (E) Ratio map of a strain bearing a donor (APY266) over a strain lacking a donor (APY358) for the left DSB end at 2 and 4 hours post-DSB induction. (F) Same as in (B) in WT (APY266), Pol3-depleted (APY1352) and Pol3– and Rfc1-depleted (APY1350) cells 4 hours post-DSB induction. (G) Left ssDNA contact profiles over a 240 kb region surrounding the donor site on chr. II in WT (APY266, n = 4), Pol3-depleted (APY1352, n = 1) and Pol3-and Rfc1-depleted (APY1350, n = 1) cells 4 hours post-DSB induction. Data smoothed over 10 kb. (H) Model for initial contacts with the donor by both NPF, homology-containing NPF disruption and loss of end-tethering. (I) Model for homology search in *S. cerevisiae*. Genome-wide homology search is (i) inhibited by static anchors, chromatin folding by cohesin, and DSB end-tethering and (ii) promoted by long-range resection and Rad51-ssDNA NPF elongation. The presence of a single long Rad51-ssDNA segment is likely sufficient to project DSB-proximal sites away from the initial chromatin environment. Two mechanisms stimulate homology search regionally: the RE, by co-opting extended NPFs. And the donor site which, owing to associations between NPFs, promote homology search in its vicinity by the end that has not yet found homology.

### Homology identification leads to NPF disruption and loss of DSB end-tethering

We previously showed that conventional Hi-C could detect homology– and Rad51-dependent DSB-donor interactions in *trans* using DSB-flanking dsDNA regions as a proxy ^25^ (**Figure 2A**, **7D**, and **S8B**). Unlike ssDNA contacts, they were not detected 2 hours post-DSB induction, and did not involve the right DSB end (**Figure 7D**). Furthermore, DSB-distal ssDNA sites interacted with the donor region as frequently as DSB-proximal sites, specifically on the homology-containing end (**Figure 7B**). These observations indicate that homology recognition leads to a collapse of the homology-containing NPF over time, bringing in proximity of the donor site the DSB-distal ssDNA sites and the dsDNA region flanking the resection front, unlike what is observed at the RE (see below, **Figure 7H)**. The absence of contacts between the donor region and DSB-distal ssDNA and dsDNA sites on the heterologous end (**Figure 7B, D, H**) suggested a loss of end-tethering occurring concurrently with the homology-containing NPF collapse. Accordingly, a partial loss of end-tethering occurred between 2 and 4 hours post-DSB induction in a donor-dependent (**Figure 7E**) and Rad51-dependent manner (**Figure S4A** and **S5A**).

### NPF collapse upon homology identification occurs prior to D-loop extension by PCNA-Pol**δ**

The kinetics of D-loop formation (2 to 4 hours) and extension (2 to 6 hours) in this strain ^32^ and the biased DSB-flanking dsDNA and DSB-distal ssDNA contacts in the direction of D-loop extension suggested that NPF collapse could occur prior to, or upon the initiation of displacement DNA synthesis by PCNA-Polδ (**Figure S8C**). To distinguish between these possibilities, we depleted AID-tagged Pol3 alone or in combination with the PCNA clamp loader subunit Rfc1 after S-phase completion (**Figure S8D**). We previously showed in these strains that this single and double depletion strategy led to a 3– and 10-fold decrease of D-loop extension, respectively ^9^. Both depletion conditions yielded highly similar results. Pol3±Rfc1 depletion did not affect G2/M-arrest, DSB resection, chromatin folding, and chromosome individualization (**Figure S8E-I**). The side-specific contacts between the left DSB-flanking dsDNA region and the donor site were retained upon Pol3±Rfc1 depletion (**Figure S8J**). Likewise, DSB-distal ssDNA sites contacted the donor region as frequently as DSB-proximal sites on the homology-containing end (**Figure 7F**). Consequently, the homology-containing NPF collapse occurs independently of PCNA-Polδ recruitment at the D-loop, and thus D-loop extension.

The asymmetric contact distribution in the direction of D-loop extension observed both for dsDNA and ssDNA sites 4 hours post-DSB induction (**Figure S8C**) was abolished upon Pol3±Rfc1 depletion (**Figure 7G** and **S8J**). It indicates that a fraction of the donor contacts originates from extended D-loops, and that D-loop extension occurs in the context of a disrupted filament (**Figure 7H**).

Finally, Pol3±Rfc1 depletion did not restore the contacts between the right dsDNA end and the donor (**Figure S8K**). Consequently, the donor-dependent loss of end-tethering results from a homology-dependent event preceding the initiation of D-loop extension, possibly upon or following DNA strand invasion (**Figure 7H**). Surprisingly, Pol3±Rfc1 depletion exacerbated the loss of DSB end-tethering observed in WT cells (**Figure S8K**). It suggests the existence of a PCNA-Polδ-dependent mechanism that holds the two ends together at the donor site, at least in our system in which homology is provided to a single end only.

## Discussion

### Regulated expansion of homology search and its determinants

We show that genomic exploration by the Rad51-ssDNA NPF is not a uniform process along the genome, the NPF, and over time. Initially local, it progressively expands to become genome-wide, a process orchestrated by independent activities imparted by chromatin folding, chromosome organization, DSB end-tethering, specialized genetic elements, long-range resection, and the extended structure of the Rad51-ssDNA NPF itself (**Figure 7I**).

Nascent, short Rad51-ssDNA NPFs sample the proximal chromatin environment. This environment is structured by static anchors (*e.g.* centromeres attachment and clustering at the SPB via microtubules) and chromatin loop extruders present in the flanking dsDNA ^25,68–71^ that expose these NPFs to a limited set of *cis* and *trans* genomic regions, and conversely restrict access to other regions (*e.g.* chromosome arm extremities). Of note, although our system does not allow to study sister chromatid interactions, the focusing of DSB-distal NPF contacts at chromatin loop base by cohesin is expected to stimulate the search by the newly resected regions on the sister chromatid first ^72^.

Long-range resection gradually provides ssDNA substrates for assembly of longer NPFs. By virtue of their rigidity ^16,73^, these NPFs progressively project DSB-proximal search units away from the initial chromatin environment. Hence, DSB-proximal and DSB-distal regions of the NPF simultaneously sample distinct genomic regions and nuclear area. This genomics-based evidence supported by modelling parallels the extended structures stretching across the nucleus recently observed in *S. cerevisiae* with a functional GFP-tagged version of Rad51 ^20^. NPF growth erodes spatial constraints and grants genome-wide homology search by DSB-proximal sections, while sections adjacent to the resection front remain engaged with the DSB-borne chromatin environment. Consistently, cohesin regulated *cis*/*trans* donor preference by DSB-proximal sequences at 2 but not 4 hours post-DSB induction, despite maintenance of cohesin-mediated loop folding ^25^.

Finally, homology search is conducted by (at least partly) paired, co-oriented NPFs, as revealed by interactions of both ssDNA ends with a region of homology to only one DSB end. This paired NPF configuration is consistent with end-tethering determined with Hi-C and cytologically with fluorescently-tagged Rad51 and other HR proteins, as well as fluorescent operator-repressor arrays in the flanking dsDNA region ^25,26,29,30,52,55^. Homology identification leads to NPF collapse specifically at the homology-containing end. This collapse is likely to limit coincidental homology identification at multiple genomic sites along a single Rad51-ssDNA NPF, and thus precludes the cascade of repeat-mediated chromosomal rearrangements initiated by multi-invasion DNA joint molecules ^6,9^.

### The possibility of genome-wide homology search lies in NPF structure

A conserved property of RecA-family proteins is to polymerize on DNA into elongated structures exhibiting persistence length orders of magnitude greater than protein-free ssDNA *in vitro* ^2^. Such structures were recently observed in bacterial and *S. cerevisiae* cells using fluorescently-tagged RecA/Rad51 proteins ^18,20,21^. Our ssHi-C data and modeling support the proposal from these cytologic studies that the possibility of genome-wide homology search lies in NPF’s atypical, rigid structure. Consequently, long Rad51-ssDNA NPFs not only serve the purpose of multiplying search units or accelerate search by inter-segmental contact sampling ^3,4^, but additionally regulates the searchable genome fraction as a function of the physical distance with the DSB-borne chromatin environment. Factors regulating the length, segmentation, rigidity, and dynamics of the NPF are thus likely homology search regulators ^20^.

Projection of NPF segments across the nuclear space entails the presence of at least one long Rad51 structure between the DSB end and the resection front. The degree of individual NPF segmentation can be directly regulated by the nucleation, growth and dissociation rates of RecA/Rad51 on ssDNA, which have been extensively studied *in vitro* ^74^. Consequently, factors such as mediators and regulators of NPF stability as well as species-specific differences in RecA/Rad51 nucleation/elongation rates likely dictate whether and with which kinetics the transition from local to genome-wide search occurs ^75–77^. The ability of RecA/Rad51 to polymerize on dsDNA^78^ may similarly achieve expanded homology search while retaining short resection tracts, a strategy that could mitigate the risk of ectopic recombination in highly repetitive genomes.

Our Gaussian polymer model consistently accounts for the observed proportion of *trans* contacts and the distribution of *cis* contacts with an apparent ssDNA persistence length of ∼20 nm. It is at least an order of magnitude higher than that of protein-free or RPA-coated ssDNA but an order of magnitude lower than that of ATP-bound RecA/Rad51-ssDNA NPF *in vitro* ^16,17,47,48,74,77,79^. This average value likely reflects a heterogeneous ensemble of ssDNA states in cells, with collapsed RPA-bound segments, contracted Rad51-ADP bound segments, and extended Rad51-ATP-bound segments co-existing on the same molecule and in varying proportion in the cell population. Consistently, Rad51-GFP structures dynamically alternate between globular and extended conformations in cells ^20^. Correcting for the fraction of cells lacking an elongated Rad51-GFP structure, our model predicts a NPF contour length of ∼1.1 μm (**Supplementary Code 1**), closely matching sizes determined cytologically ^20^.

### Long-range resection promotes genome-wide homology search

Long-range resection primarily conducted by Exo1 progresses unabated at ∼4 kb/h in mitotically-dividing *S. cerevisiae* cells ^80,81^. Such extensive generation of ssDNA largely exceeds the length of homology needed for efficient ectopic repair by HR ^82–84^, which questioned the function of long-range resection ^65^. Here, we show that Exo1 promotes genome-wide homology search by DSB-proximal ssDNA. It likely represents and adaptation of the search strategy in the event of a failure of swift repair using spatially favored chromatin (*e.g.* the sister chromatid) ^85^. Consistently, long-range resection is required only when the donor is provided >50 kb away from the DSB site or on another chromosome ^65^, *i.e.* outside the typical range of high ssDNA contacts determined here in WT cells early post-DSB induction and in *exo1* mutants.

Since as little as 2 kb of ssDNA is sufficient to assemble a 1 μm-long NPF (*i.e.* the diameter of a yeast nucleus), ssDNA is unlikely to be limiting for extended NPF assembly, even in the absence of Exo1. Consequently, and as discussed above, we favor the view that most NPF segments are short, and that the excess of ssDNA produced by long-range resection increases the probability of assembling at least one long (*i.e.* >1 kb ∼ 500 nm) segment (**Figure 7I**).

### Role of DSB end-tethering in coordinating homology search by the two NPFs

Maintenance of end-tethering involves several handovers over the course of DSB repair: initially mediated by the Mre1-Rad50-Xrs2^NBS1^ (MRX^N^) complex prior to and upon short-range resection, it is followed by Exo1 and the Ddc1-Mec3-Rad17 checkpoint clamp (human 9-1-1 clamp) independently of its checkpoint function ^25,27,55^. This tethering at the level of the resection front is required for coordinated search by both NPFs. Such coordination is supported by the predominance of single Rad51-GFP structures over branched and complex structures observed cytologically ^20^.

Pre-association between DSB ends facilitates SDSA repair steps downstream of D-loop extension ^86,87^. Here we show that end-tethering promotes coordinated homology search between the two ends: identification of homology at the terminus of one end stimulates homology search in that region by the terminus of the opposite end. Such enhancement of local homology search conditional to identification of homology by one of the two ends likely promotes concurrent donor identification and/or invasion by both ends, a possible precursor of double Holliday junction formation ^88^.

Intriguingly, in the absence of homology to the second end, end-tethering is progressively lost, coincidentally with a disruption or collapse of the homology-containing Rad51-ssDNA NPF. This observation extends the sequence of handovers required for the maintenance of DSB end-tethering taking place along the HR pathway at the synaptic step upon homology identification of the same donor site by both DSB ends. Absence of shared homology is likely to signal an ectopic invasion event, and loss of end-tethering in this context may inhibit the formation of repeat-mediated rearrangements. PCNA-Polδ also promoted end-tethering in our system lacking homology to the second end. The relevance and mechanism of this association remain to be determined.

### Not all genomic sites are equal for homology search

We show that the genome is not equally sampled during homology search, and that multiple mechanisms exist to enhance it at specific genomic loci, conditionally or constitutively: cohesin promotes sampling of regions at the base of chromatin loops (and presumably the sister chromatid) by ssDNA sites newly generated by resection; centromere clustering promotes search on other centromeres; and DSB end-tethering enhances homology search around the donor site if homology has been identified by one of the two end. We identified a fourth mechanism specific of chromosome III that engages pioneer, extended NPFs. It relies on the Recombination Enhancer (RE), an autonomous genetic element involved in biasing donor usage in mating-type switching or ectopic repair in *MAT***a** cells ^60,62,67^. Our observations strongly suggest that RE stimulates recombination with nearby donor by engaging the NPF, which locally enhances homology search.

The RE function depends on its binding by multiple copies of the conserved fork-head-family transcription factor Fkh1 ^62,89,90^. It raises the possibility that Fkh1, present as a monomer at gene promoters throughout the genome ^91,92^, may similarly attract the NPF and locally bias genome-wide homology search. By seeding homology sampling events regularly, such mechanism may drastically reduce search complexity and could be harnessed to inhibit repeat-mediated rearrangements by occluding repeated elements. PRDM9 has been proposed to assume a similar function downstream of DSB formation during mouse meiosis ^93^.

### Study of other DNA metabolic processes with ssHi-C

ssDNA is a hallmark of various fundamental DNA metabolic processes. By directly targeting ssDNA, ssHi-C enables for the first time the site-specific determination of the contacts made by DNA while it is being involved in such processes, even if only a subpopulation of cells exhibits ssDNA at the site of interest. Since ssHi-C operates according to Watson-Crick base-pairing rules and is protein-independent, it can in theory be applied in any organism for which the genome sequence is known, without a need for specific reagents (*e.g.* antibody) or genetic engineering. Consequently, ssHi-C is a universal method for the study of the spatial aspects of DNA metabolic processes such as DNA replication, mitotic and meiotic recombination, bacterial conjugation, transposition and the replication of certain types of viruses.

## Acknowledgements

We are grateful for helpful discussions with Piazza, Jost and Koszul lab members. We thank Wolf-Dietrich Heyer and Angela Taddei for their critical reading of the manuscript. We thank Romain Koszul for critical reading of the manuscript and for sharing expertise and protocols. We are grateful to Karine Dubrana for sharing unpublished data regarding intra-chromosomal distances in G2/M cells.

This research was supported by the CNRS as part of its Momentum program and the European Research Council (ERC) under the European Union’s Horizon 2020 to A.P. (ERC grant agreement 851006) and the Agence Nationale de la Recherche to DJ (ANR-21-CE13-0037-02, ANR-22-CE12-0035-03).

## Author contributions

Conceptualization: AP; Experiments: AD, JS, AT, PM, AP; Data analysis: NM, AP; Software development: NM, LA, LM; Modeling: DJ; Data interpretation: AP, DJ; Supervision: AP, LM; Funding Acquisition: AP; Writing: AP.

## Methods

### Saccharomyces cerevisiae strains

Genotypes of *Saccharomyces cerevisiae* (W303 *RAD5*+ background) strains are listed in **Table S1**. The inducible *HO* expression construct *pGAL1-HO* at *trp1* (chr. IV), the mutagenesis of the endogenous HO cut-site at *MAT* (chr. III), the DSB-inducible construct replacing *URA3* (chr. V) and the donor sequence replacing *LYS2* (chr. II) have been described previously ^31,32^. The DSB-inducible construct contains 327 bp of a fragment of the PhiX phage genome flanked by multiple restriction sites (including *DpnII*) making a unique 453 bp sequence pad for annealing of oligonucleotides in ssHi-C, a 2,086 bp-long sequence corresponding to the first half of the *LYS2* gene (+4 to +2,090), and a 117 bp HO cut-site. It is positioned at position 116,167 (S288c coordinates), replacing *URA3*. A 2,086 bp-long ectopic donor (+4 to +2,090 of the *LYS2* gene) is present at its endogenous locus on chr. II.

The construct *his3::pADH1-OsTIR1-9Myc::HIS3* for OsTir1 E3-Ubiquitin ligase expression, as well as the Scc1-(Pk3-)AID, Scc1-6HA and Cdc45-(FlagX5-)AID constructs have been described previously ^32,57^. The *rad51::kanMX, exo1::kanMX*, and *exo1-D173A* mutations were previously described ^25^. The *RFC1-AID-9Myc::hphMX* construct has been described previously ^95^. The *POL3-iAID* construct has been described previously ^96,97^. The constructs for *TIR1* and *TetR* expression integrated at *SSN6* have been described previously ^97^. Seamless deletion of the RE on chr. III (R64-2-1 coordinates chrIII:29107-30801) was achieved upon templated CRISPR/Cas9-induced editing ^98^ (**Table S2**). Insertion of the 700 bp RE at position 845464 on chr. IV was achieved by transformation of a strain containing *URA3* at that site with a PCR product amplified with primers RE-atchrIV-845464-F/R (**Table S2**) and selecting for 5-FOA-resistant colonies. Gene deletions, fusions and point mutations were verified by PCR and/or Sanger sequencing, as well as from bam sequence files generated upon alignment of Hi-C or ssHi-C reads. Protein tagging was verified by Western blotting.

### Culture media

Liquid YPD (1% yeast extract, 2% peptone, 2% dextrose) and YEP-lactate (1% yeast extract, 2% peptone, 2% lactate) were prepared according with standard protocols ^99^.

### Induction of DSB formation in cells synchronously released in S-phase

Exponentially growing cultures in YEP-lactate (30°C) were synchronized at the G1/S transition by addition of 1 µg/ml alpha-factor (GeneCust) every 30 minutes for 3 hours. Cells were washed 3 times with 50 mL of pre-warmed YEP-lactate and released in S-phase in YEP-lactate supplemented with 2% galactose to simultaneously induce HO expression. The cell-cycle stages were verified by flow cytometry.

### Protein depletion

Crosslinked pellets originating from time courses performed in ref. ^25^ were reprocessed for ssHi-C. Protein depletion and flow cytometry analysis were reported in ref. ^25^. Briefly, Cdc45-(FlagX5-)AID depletion was induced in cells synchronized with alpha-factor by addition of 2 mM 3-Indole Acetic Acid (IAA, *i.e.* Auxin) solubilized in DMSO, or of the equivalent DMSO concentration (0.35% final) as control, 1 hour before release from the G1 arrest.

Scc1-(Pk3-)AID depletion was induced at the time of release in S phase by addition of 2 mM IAA (or the equivalent DMSO concentration), together with DSB induction. Proteins levels were determined by Western blot.

Inhibition of *POL3-iAID* expression and depletion of Pol3-iAID independently or together with Rfc1-AID-9Myc was conducted similarly to ^9,96^. Transcriptional inhibition was induced upon 50 μg/mL doxycycline and protein depletion induced upon 1.5 mM IAA addition 1 hour after S-phase release and HO induction, to allow for bulk DNA replication to take place.

### Flow cytometry

Approximately 107 cells were collected by centrifugation, re-suspended in 70% ethanol and fixed at 4°C for at least 24 hours. Cells were pelleted, resuspended in 1 mL of 50mM Sodium citrate pH 7.0 and sonicated 10 seconds on a Bioruptor. After washing, cells were treated with 200 µg of RNase A (Euromedex, cat.9707-C) at 37°C overnight. Cells were then washed and incubated for 30 minutes with 1 mL of 50 mM Sodium citrate pH 7.0 with 16 µg of Propidium iodide (Fisher Scientific, 11425392). Flow cytometry profiles were obtained on a MACSQuant machine and analyzed using Flowing Software 2.5.1.

### Hi-C and ssHi-C procedures

- Annealing and capture oligonucleotide design

Annealing oligonucleotides have been designed according to the following rules:

○ Be 80 nt-long.
○ Overlap a DpnII restriction site and a MfeI or SspI restriction site located at less than 40 bp from the DpnII site.
○ Asymmetrically positioned relative to the DpnII site so has to ensure sufficient base-pairing near the DpnII site (> 4 bp) and maximize the length of annealed dsDNA produced on the MfeI/SspI-containing side.
○ Contain 4-6 single nucleotide substitutions introduced as evenly as possible within the MfeI/SspI-containg side. One of these substitutions must inactivate the MfeI/SspI site. Other substitutions are chosen preferentially to inactivate cryptic DpnII (*e.g.* <u>A</u>ATC instead of GATC) and HinfI (*e.g.* <u>A</u>ANTC instead of GANTC). Substitutions are not introduced within 4 bp of the DpnII site. Finally, they must be as evenly distributed as possible.
○ Oriented so as to anneal to the ssDNA present in the cell.
○ Be HPLC-or PAGE-purified.
○ Have their 5’ and 3’ ends chemically blocked to prevent artefactual biotinylation and ligation in the absence of annealing to, and digestion of, their DpnII target site. Blockers used are 5-methyl-deoxycytidine at the 5’ end and the phosphoramidite C3-spacer at the 3’ end.

Capture oligonucleotides target the dsDNA part restored by the annealing oligonucleotides, apart from control dsDNA capture oligonucleotides that target single dsDNA fragments without specific constrains. They contain a 5’ Biotin. Annealing and capture oligonucleotides were synthesized by Integrated DNA technologies. Annealing and capture oligonucleotides sequences are provided in **Table S2**. For an example of the design of the annealing and capture oligonucleotide for the DpnII site L2599 (corresponding to the restriction fragment 18614), see **Figure S1B**.

- Cell fixation with formaldehyde and EGS

Cell fixation is adapted from Micro-C-XL ^100^. Approximately 1.5×10^9^ haploid cells were fixed with 3% formaldehyde (Sigma-Aldrich, cat. F8775) for 30 minutes at RT at 120 rpm. Formaldehyde was quenched with 330 mM glycine for 20 minutes at RT at 120 rpm. Cells were washed twice with cold water at 3,000 g for 10 minutes. Pellets were split in two tubes and frozen at –80° C.

Cell pellets were thawed in ice, resuspended in 10 mL of Zymolyase solution (0.25 mg/mL Zymolyaze 100T (Carl Roth, cat. 9329), 1 M D-sorbitol (Sigma-Aldrich, cat. S6021), 50 mM Tris-HCl pH 7.5, 10 mM 2−mercaptoethanol (Sigma-Aldrich, cat. M3148)) and spheroplasted at 210 rpm for 18 minutes at 30° C (cells spheroplasting is checked under the microscope and the incubation time adjusted accordingly). Spheroplasts are pelleted at 3,000 g for 10 minutes at 4° C, washed with 4 mL cold 1X PBS, and resuspended in a fresh EGS crosslinking solution (3 mM ethylene glycol bis(succinimidyl succinate) (Fisher Scientific, cat. 10350924) in 1X PBS), and incubated at 30° C for 40 minutes at 100 rpm. EGS was quenched with 400 mM glycine at RT for 5 minutes at 100 rpm. The crosslinked cellular material was pelleted at 3,000 g for 10 minutes at 4° C and washed twice in cold 1X PBS supplemented with anti-protease (Sigma-Aldrich, cat. 11836170001). Pellets were snap-freezed in liquid nitrogen and stored at –80°C.

- Hi-C procedure with restriction site restoration in ssDNA

The crosslinked pellets were thawed in ice and resuspended in cold H_2_O supplemented with anti-protease. Between 2 and 5.10^7^ cells were used for the remainder of the procedure. The ssHi-C procedure uses the Arima Hi-C kit (Arima Genomics, cat. A410079), which employs a dual restriction digestion (DpnII and HinfI) yielding a median fragment length of 108 bp in *S. cerevisiae.* The protocol follows manufacturer’s instructions with the following modification: after the lysis step, the pool of HPLC-or PAGE-purified annealing oligonucleotides (stock concentration 100 nM each) is added at a final concentration of 7 nM each. Annealing occur during the subsequent incubation at 62° C for 10 minutes.

- Sequencing library preparation

DNA was fragmented into 300-400 bp fragments using Covaris M220 sonicator. Preparation of the libraries for paired-end sequencing on an Illumina platform was performed using the Thermofisher Collibri ES DNA Library Prep Kit for Illumina Systems with UD indexes (cat. A38606024) following manufacturer’s instructions. The library was amplified in triplicate PCR reactions using oligonucleotides corresponding to the Illumina sequence adaptors (5’-AATGATACGGCGACCACCGAGATCTACAC-3’ and 5’-

CAAGCAGAAGACGGCATACGAGAT-3’) and Phusion DNA polymerase (New England Biolabs,

cat. M0531) for 11 cycles. PCR products were purified with AMPure XP beads (Beckman-Coulter, cat. A63881) and resuspended in pure H_2_O. The Hi-C library is quantified using the Qubit DNA high sensitivity kit (Thermo Scientific, cat. Q32851) on a Qubit 2 fluorometer (Thermo Scientific, cat. Q32866).

- ssHi-C library Capture

The ssHi-C library consists in an enrichment of ssDNA contacts from the Hi-C library using a panel of biotinylated “capture” oligonucleotides. 500 ng of PCR products were saturated with 2 μg of unlabeled Cot1 human DNA (Invitrogen, cat. 15279-011) and 0.2 nmol of oligonucleotides corresponding to the universal Illumina adaptors (see above) and dehydrated at 60° C for 1 hour in a Speedvac (Thermo Scientific, cat. DNA130-230). DNA was resuspended in 1X xGen Hybridization buffer and 0.15X of xGen Hybridization Enhancer (Integrated DNA Technologies, cat. 10010352), and the pool of biotinylated capture oligonucleotides (stock 1.8 μm each) at a final concentration of 0.6 μM. DNA is denatured at 95° C for 5 minutes, and annealing occurs at 65° C for 4 hours. Biotin pulldown on streptavidin-coated dynabeads C1 (Invitrogen, cat. 65001) was performed according to manufacturer’s instructions, and DNA was recovered in pure H_2_O. PCR amplification (10 cycles) and DNA purification on AMPure XP beads was carried out as before. The capture procedure was repeated once and the final library was amplified for only 8 cycles.

- Optional: elimination of the dsDNA-borne contacts

This step eliminates the chimeric molecules involving the fragments of interest (*i.e.* targeted by annealing and capture oligonucleotides) that were in a dsDNA form in the cell population at the time of crosslink, upon digestion by restriction enzymes whose site was eliminated in the annealing oligonucleotide (rationale **Figure S1A**) 300 ng of PCR products were digested at 37° C for 1 hour with 10 units of MfeI-HF and SspI-HF in 1X rCutsmart buffer (New England Biolabs, cat. R3589, R3132 and B7204, respectively) and the enzymes heat-inactivated at 65° C for 20 minutes. DNA was purified with AMPure XP beads, resuspended in pure H_2_0, and quantified using the Qubit DNA high sensitivity kit (Thermo Scientific, cat. Q32851) on a Qubit 2 fluorometer (Thermo Scientific, cat. Q32866).

- Paired-end illumine sequencing

ssHi-C libraries were paired-end sequenced (2×150 bp) on an Illumina Novaseq6000 platform by Novogene UK.

### Protein extraction and western blotting

Protein extracts for western blot were prepared from 5.10^7^ to 10^8^ cells. Cells were lysed in cold NaOH buffer (1.85 N NaOH, 7.5% v/v beta-Mercaptoethanol) for 10 min in ice. Addition of trichloroacetic acid (15% final) for 10 min in ice allowed protein precipitation. After centrifugation at 15,000 g for 5 min, the pellets were resuspended in 100 µL of SB++ buffer (180 mM Tris-HCl pH 6.8, 6.7 M Urea, 4.2% SDS, 80 µM EDTA, 1.5% v/v Beta-mercaptoethanol, 12.5 µM Bromophenol blue). Denaturation was performed by heating 5 min at 65°C. Pre-cleared extracts were resolved on 12% precast polyacrylamide gel (Bio-Rad, cat. 4561043) and blotted on a PVDF membrane (GE Healthcare, cat. 10600023). Membranes were probed with mouse anti-AID antibody diluted at 1:1000 (CliniSciences, M214-3), or anti-GAPDH antibody diluted at 1:10000 (Invitrogen, MA515738), and revealed with an HRP-conjugated anti-mouse IgG antibody diluted at 1:10000 (Abcam, ab6789) using Immobilon Forte western HRP substrate (Merck, WBLUF0100) and a Chemidoc MP Imaging system (BioRad).

## Data analysis

### Hi-C read alignment and generation of contact maps

Paired-end reads are aligned and processed using Hicstuff *pipeline* function in “cutsite” mode ^101^. Briefly, pairs of reads are aligned iteratively and independently using Bowtie2 in its most sensitive mode to the *S. cerevisiae* reference genome deleted for the *HML, MAT, HMR, URA3,* and *LYS2* sequences. Each uniquely mapped read was assigned to a restriction fragment. Uncuts, loops and circularization events were filtered as described in ^102^ and PCR duplicates discarded.

### ssHi-C read alignment

The ssHi-C alignement pipeline is depicted in **Figure S1B**. First, the reference genome is edited so as to introduce the sequence of the annealing oligonucleotides (or its reverse-complement) in place of the endogenous matching sequence on the Watson strand. The endogenous sequences are concatenated as an artificial “trash” chromosome. This modified genome sequence containing both the original reference and the modified sequence is used for competitive mapping of the reads. Reads originating from ligation events made by the fragment in a ssDNA form will map at the SNP-containing fragment, while reads originating from ligation events made by the fragment in a dsDNA form will map on the unmodified sequence on the “trash” chromosome. Read mapping is conducted as for Hi-C using the Hicstuff *pipeline* function in “cutsite” mode and with a quality threshold of 20. A custom ssHi-C pipeline further isolate contacts made by ssDNA fragments to generate “4C-like” plots, compute statistics, ponder contacts using an external reference, and determine the absolute number of unique contacts per fragment.

### Generation of ratio maps

Ratio maps were generated from sparse matrices using Hicstuff *view* function, normalized (SCN normalization) ^102^, log-transformed and binned. Alternatively, ratio maps were generated with Serpentine ^94^. Briefly, two matrices binned at 1 kb of the chromosome of interest were subsampled to contain the same number of contacts. Comparison of contact maps was performed using the default threshold parameters (50 and 5) over 10 cycles, and the detrending constant set to 0.

### Generation of aggregated contact maps

Intra-and inter-chromosomal centromere pile-up contact maps were generated with Chromosight *quantify* with default parameters ^103^.

### Computation of the contact probability as a function of genomic distance

Computation of the contact probability as a function of genomic distance P_c_(s) and its derivative have been determined using Hicstuff *distance law* function^9^ with default parameters, averaging the contact data of entire chromosome arms.

### Quantification of coverage from Hi-C data

Quantification of coverage from Hi-C data was determined with custom scripts from sparse contact matrices generated with Hicstuff *pipeline* function. Briefly, the sum of total contact frequencies within 1 kb bins was computed along the broken chromosome for each sample, and normalized onto the average coverage of undamaged cells obtained from a biological replicate.

### Statistics

The proportion of intra-chromosomal contacts were compared using a non-parametric Mann-Whitney Wilcoxon rank sum test, two-tailed. Statistical cut-off was set at 0.05. P-values below this cut-off are denoted by a star (*) in figures. Statistical tests were performed under R 3.6.2. Linear regression and data smoothing was performed under Graphpad Prism 9. No statistical method was used to predetermine sample size. No data were excluded from the analyses. The experiments were not randomized. The investigators were not blinded to allocation during experiments and outcome assessment.

### Gaussian contact probability model

Briefly, we modeled the NPF as a homogeneous semi-flexible chain characterized by a persistence length *L_p_* and a contour length *L_c_*. *L_p_* corresponds to the typical lineic distance along the chain below which the NPF could be considered as rigid, and *L_c_* to the lineic distance along the chain between the resection front and the DSB. Since the overall genome organization is not significantly affected during DNA repair compared to normal G2/M, we posited that the probability for a ssDNA probe to contact a dsDNA fragment depends on the relative 3D position of the resection front compared to the dsDNA locus in normal condition (no break) and on the probability of the ssDNA probe inside the NPF emanating from the resection front to be found in close proximity to the dsDNA. The NPF thus plays the role of a fishing rod in the hands of a fisherman located at the resection front with a hook attached at the probe position (**Figure 2H** of the main text). Assuming that the distribution of 3D relative positions between any pairs of loci to be Gaussian^104^, we could then express the average contact probability between any ssDNA probe with any dsDNA fragment as a function of the contact probability in normal conditions, of the statistics of position of the resection front and of the spatial organization of the NPF (see **Supplementary Code 1**). The contact probability in normal conditions is given by Hi-C experiments. The statistics of resection front is extracted from a simple 1D stochastic model of the resection front around the DSB and fitted on the experimental Hi-C coverage data in DSB conditions. The spatial properties of the NPF are given assuming a worm-like-chain statistics depending on *L_p_* and *L_c_*. The only unknown parameter in the model is *L_p_* that we then varied to fit the experimentally-measured fraction of *trans* contact. All the mathematical details of the model are given in the **Supplementary Code 1**.

## Data availability statement

Data will be made available upon publication.

## Software availability

Hicstuff ^101^ (version 3.2.1 available at https://github.com/baudrly/hicstuff)

Chromosight ^105^ (version 1.6.3, available at https://github.com/koszullab/chromosight)

Serpentine ^94^ (version 0.1.3, available at https://github.com/koszullab/serpentine)

Tinycov (version 0.3.1, available at https://github.com/cmdoret/tinycov)

TinyMapper (version 0.1, available at https://github.com/js2264/tinyMapper)

R (version x64 3.6.2 is available online at https://cloud.r-project.org/)

Bowtie2 ^106^ (version 2.3.5.1 available online at http://bowtie-bio.sourceforge.net/bowtie2/)

Samtools ^107^ (version 1.3.1 available online at https://github.com/samtools/samtools)

Bedtools ^108^ (version 2.27.1 available online at https://github.com/arq5x/bedtools2)

Cooler ^109^ (version 0.9.1 available online at https://github.com/open2c/cooler)

Flowing Software (version 2.5.1 available online at https://flowingsoftware.com/download/)

