## Supplementary information for "Mechanism of homology search expansion during recombinational DNA break repair"

### Supplementary figures and table legends

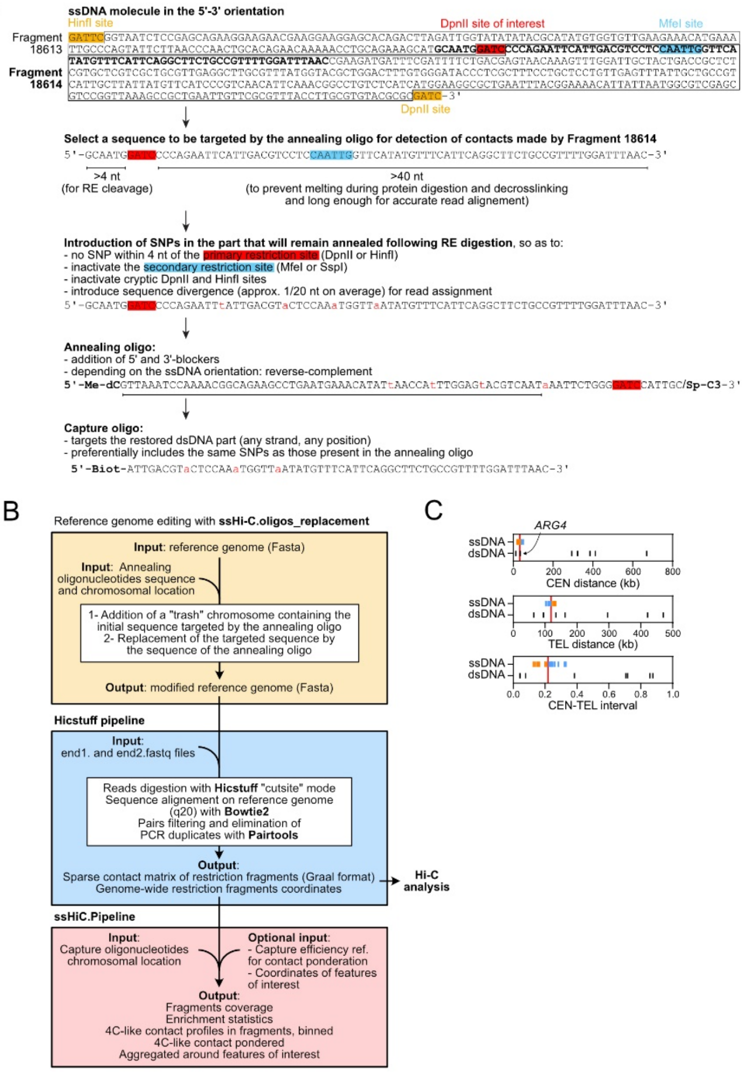


**Figure S1: Rationale of ssHi-C, oligonucleotides design and reads assignment. (Related to Figure 1)**

(A) Example of annealing and capture oligonucleotide design for a DpnII site located 2599 bp away from the HO cut-site (L2599 in **Figure 1E**).

(B) Pipeline for alignment of ssHi-C reads and filtering and analysis of ssDNA contacts. The reference genome is modified so as to introduce the sequence of the annealing oligonucleotides. This modified genome sequence containing both the original reference and the modified sequence is used for competitive mapping of the reads. Reads originating from ligation events made by the fragment in a ssDNA form will map at the SNP-containing fragment, while reads originating from ligation events made by the fragment in a dsDNA form will map on the unmodified sequence. In the case of MfeI/SspI digestion, no reads map at the unmodified sequences (see **Figure S2B**).

(C) Location of the captured ssDNA and dsDNA sites relative to the centromere and telomere. The *ARG4* dsDNA control is located at an equivalent distance from a centromere as the ssDNA sites.

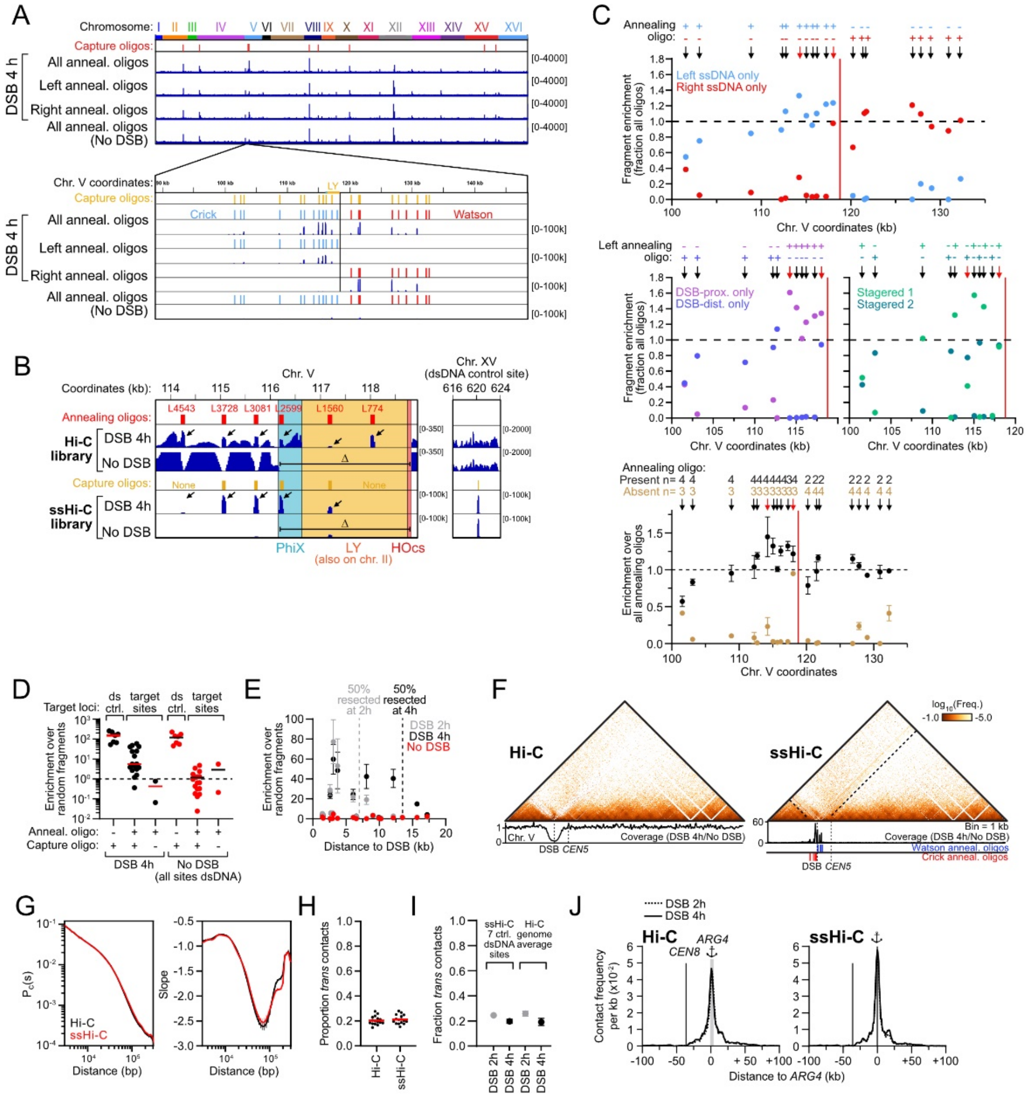

**Figure S2: ssHi-C detects both ssDNA and dsDNA contacts. (Related to Figure 1)**

(A) Representative read coverage of ssHi-C libraries obtained in WT cells (APY266) 4h post-DSB induction with annealing oligo mixes targeting ssDNA either on both sides of the DSB (All) or only the left or right ends. A ssHi-C library containing all annealing oligonucleotides but lacking endogenous ssDNA (No DSB; APY142) does not exhibit read enrichment at the target sites. The capture oligonucleotide mix was kept constant in all conditions.

(B) Left: Representative Hi-C and ssHi-C results in WT cells (APY266) 4h post-DSB induction, or in cells lacking the DSB-inducible construct. Restoration of restriction sites is DSB-dependent, and their enrichment depends on the presence of capture oligonucleotides. ssDNA sites restoration is partial and varies in efficiency from site to site (black arrows). Right: A representative control dsDNA site on chr. XV is efficiently captured with and without DSB.

(C) Targeted fragment enrichment in annealing oligonucleotide dropout experiments relative to ssHi-C library containing all annealing oligonucleotides. The bottom plot is the average of data in the upper plots.

(D) dsDNA and ssDNA sites enrichment over genome average in the presence and absence of a DSB. Each dot represents an independent site.

(E) ssDNA sites enrichment over genome average at a distance from the DSB site at 2 and 4 hours post-DSB induction, or in the absence of DSB. The extent of resection partly accounts for the heterogeneity of ssDNA enrichment.

(F) Hi-C and ssHi-C contact maps of chr. V in WT cells 4 hours post-DSB induction, revealing ssDNA-specific contact stripes emanating from the DSB region and spreading along the chromosome. Same sample as in **Figure 1F**. Hi-C and ssHi-C coverage is shown. Bin: 1 kb.

(G) Comparison of the $P_c(s)$ and its derivative obtained from Hi-C and corresponding ssHi-C libraries of WT cells 4 hours post-DSB induction. Data represent mean ± SEM of 4 biological replicates.

(H) Comparison of the *trans* contacts for each chromosome obtained from Hi-C and corresponding ssHi-C libraries of WT cells 4 hours post-DSB induction. Data are average from 4 biological replicates. Red bar: median.

(I) Comparison of proportion of *trans* contacts determined in WT cells 2 and 4 hours post DSB-induction at the 7 captured dsDNA sites (ssHi-C, left) or averaged over the genome (Hi-C, right). Data represent mean ± SEM of 4 biological replicates.

(J) 4C-like representation of the distribution of contacts by the *ARG4* locus before capture enrichment (Hi-C; left) and after capture enrichment (ssHi-C; right). Data smoothed over 4 kb.

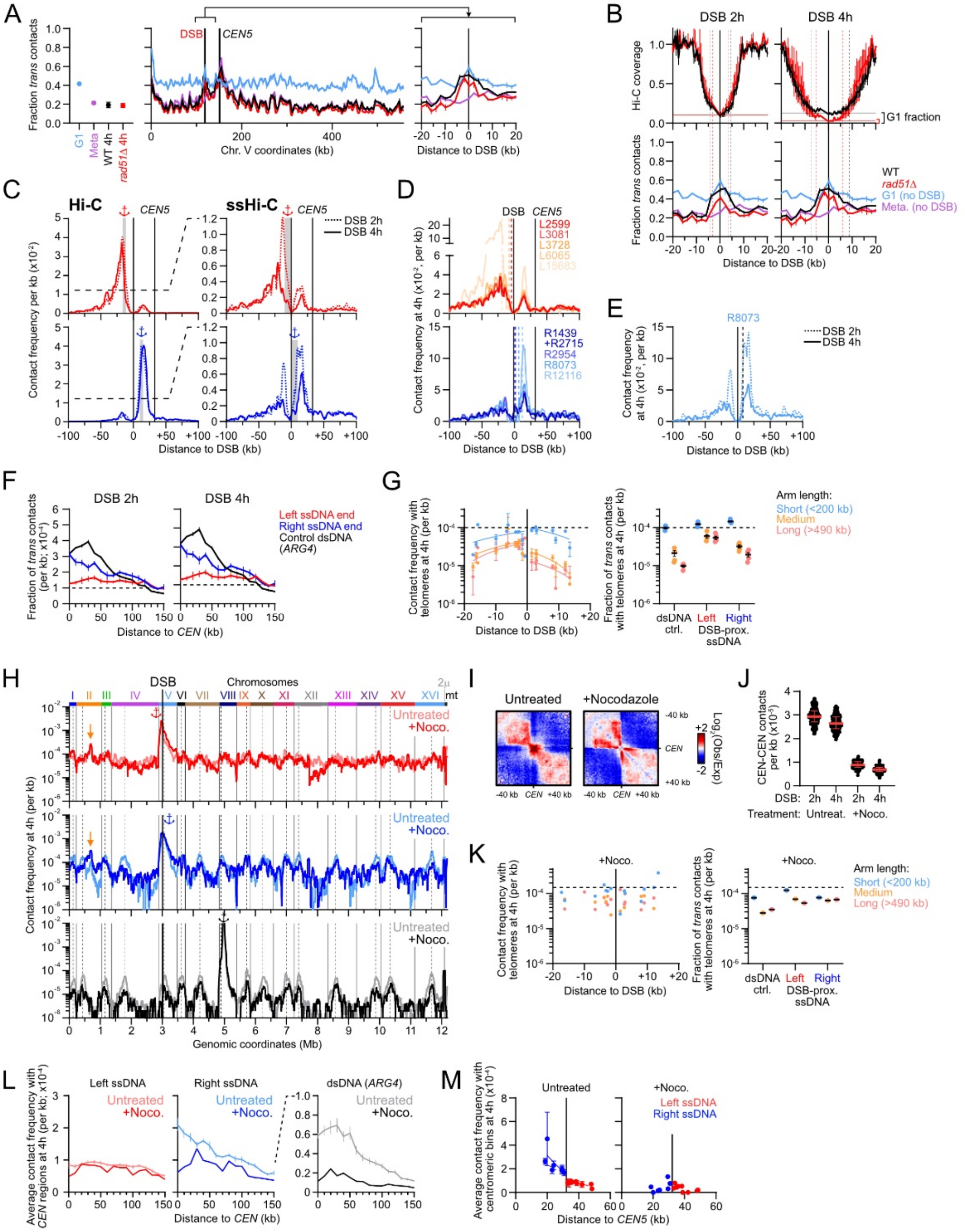


**Figure S3: ssDNA contacts are constrained by nearby centromere anchoring at the SPB, but to a lower extent than DSB-flanking dsDNA sites. (Related to Figure 2).**

(A) Proportion of *trans* contacts determined by Hi-C along chr. V (right) and averaged over the genome (left) in WT cells without DSB arrested for 4 hours in G1 with alpha-factor and in metaphase upon repression of *CDC20* expression [1], and in WT and *rad51Δ* cells 4 hours post-DSB induction (n = 2, 2, 4, and 2 biological replicates in APY266, APY537, APY266 and APY679, respectively). A partly

Rad51-independent increase in *trans* contacts is observed in the ~30 kb DSB-surrounding region, to levels observed in undamaged G1 cells.

(B) Same data as in (A) with the addition of *trans* contact profiles at 2 hours post-DSB induction, and Hi-C coverage in WT and *rad51Δ* cells. The region exhibiting Rad51-independent increase in *trans* contacts extends over to the lowest Hi-C coverage section (~10 kb at 4 hours), which corresponds to a flat unresected region. Reads in this unresected regions likely originates from a minor fraction of the cell population (<5 %) still in G1 (**Figure 1B**). Consequently, Hi-C contact data in the ~10 kb surrounding the DSB region are dominated by this unresected G1 sub-population. It explains why the proportion of *trans* contacts is identical to a G1-arrested population, and independent on Rad51.

(C) Zoom of the Hi-C and ssHi-C data in **Figure 2A** around the DSB region. Data smoothed over 4 kb.

(D) Contact distribution in the DSB region for individual ssDNA sites. Data from the same samples as in D, smoothed over 4 kb.

(E) Contact distribution in the DSB region for the L8073 ssDNA site, located in the vicinity of the resection front at 2 hours, and at ~8 kb from the resection front at 4 hours post-DSB induction.

(F) Frequency of *trans* contacts with centromere-surrounding regions for the left and right DSB ends and the control (*ARG4*) dsDNA site in WT cells 2 and 4 hours post-DSB induction, determined by Hi-C and ssHi-C. The Hi-C viewpoint corresponds to a 5 kb bin located 5-10 kb away from the DSB site. Data represent mean ± SEM of 4 biological replicates.

(G) Among *trans* contacts only, average contact frequency between ssDNA sites and 30 kb telomeric regions as a function of chromosome arm length in WT cells. Data show mean ± SEM of 4 biological replicates. Linear regressions for the left and right ssDNA sites are shown.

(H) Average genome-wide contact distribution of ssDNA sites on the left and right of the DSB, and for a control dsDNA site located at the same distance from a centromere as the DSB site 4h post-DSB induction in untreated (n = 4) or nocodazole-treated (n = 1) WT cells (APY266). Anchors: viewpoints. Orange arrow: donor. Dotted lines: centromeres. Data smoothed over 30 kb.

(I) Aggregated ratio maps of 29 kb windows centered on the 16 centromeres over randomly chosen positions on the diagonal, from Hi-C data 4 hours post-DSB induction in untreated and nocodazole-treated WT cells.

(J) Contact frequency between all pairs of centromeres, from Hi-C data 2 and 4 hours post-DSB induction in untreated and nocodazole-treated WT cells. Median and interquartile range is shown.

(K) Same as (G) for nocodazole-treated WT cells.

(L) Average contact frequency with *trans* centromere-surrounding regions for the left and right ssDNA ends and the control (*ARG4*) dsDNA site in 4 hours post-DSB induction. From data in (H).

(M) Average contact frequency with *trans* centromeres (1 kb bin) for each ssDNA sites, as a function of the distance to the DSB sites. From data in (H). Error bars show SEM.

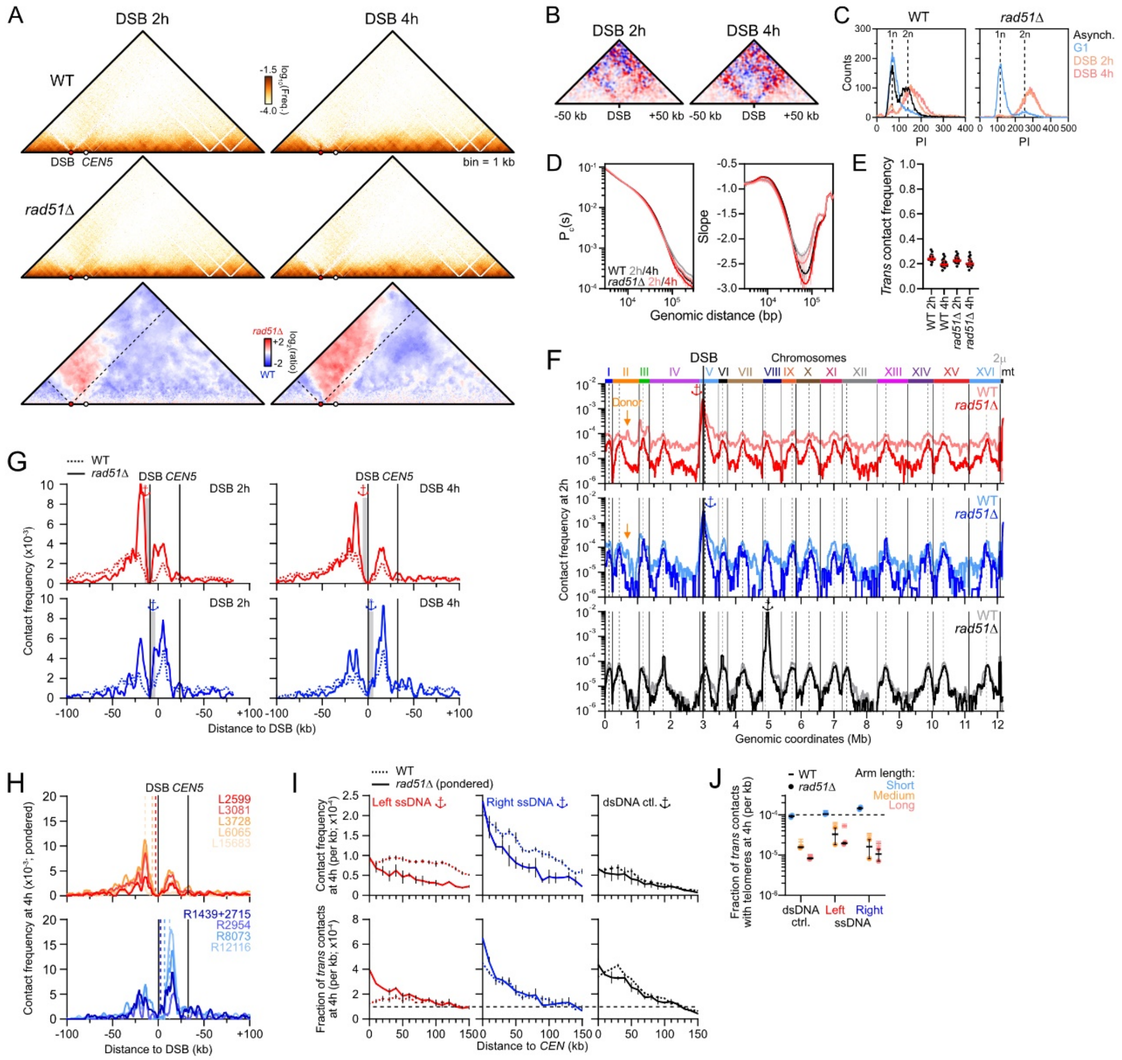

**Figure S4: Rad51 promotes genome-wide homology search by DSB-proximal ssDNA sites. (related to Figure 3)**

(A) Representative Hi-C contact maps and Serpentine ratio maps of chr. V in WT (APY266, n = 4) and *rad51Δ* (APY679) cells 2 and 4 hours post-DSB induction.

(B) Ratio maps of the 100 kb region surrounding the DSB site of the data in (A). Bin: 2 kb.

(C) FACS profiles of WT (APY266) and *rad51Δ* cells (APY679).

(D) $P_c(s)$ and derivatives of WT and *rad51Δ* Hi-C data in (A). Data: mean ± SEM of 4 and 2 biological replicates, respectively.

(E) *Trans* contact frequency determined from WT and *rad51Δ* Hi-C data (n =4 and 2 biological replicates, respectively). Each dot represents a chromosome. Bar: Median. Error bars: interquartile range.

(F) Average genome-wide contact distribution of the left and right DSB ends 2 hours post-DSB induction in WT cells (APY266) and *rad51Δ* cells (APY679; pondered). Other legends as in **Figure 2A**.

(G) Zoom of the ssHi-C data in (F) around the DSB region, smoothed over 4 kb.

(H) Pondered contact distribution in the DSB region for individual ssDNA sites on the left (top) and right (bottom) DSB ends. Data from the same samples as in (F-G), smoothed over 4 kb. Dotted lines represent the position of individual viewpoints.

(I) Contact frequency of the average left ssDNA, right ssDNA, and of a control dsDNA site (*ARG4*) with *trans* centromere-flanking regions of data in **Figure 3B**. Top: frequency of total contacts. Bottom: Fraction of *trans* contacts only. Data: mean ± SEM. Dotted line: genome average.

(J) Among *trans* contacts only, average contact frequency between DSB-proximal ssDNA sites and 30 kb telomeric regions as a function of chromosome arm length in *rad51Δ* cells 4 hours post-DSB induction. Data show mean ± SEM of 2 biological replicates. Dotted line indicates the genome average. WT data are from **Figure S3J**.

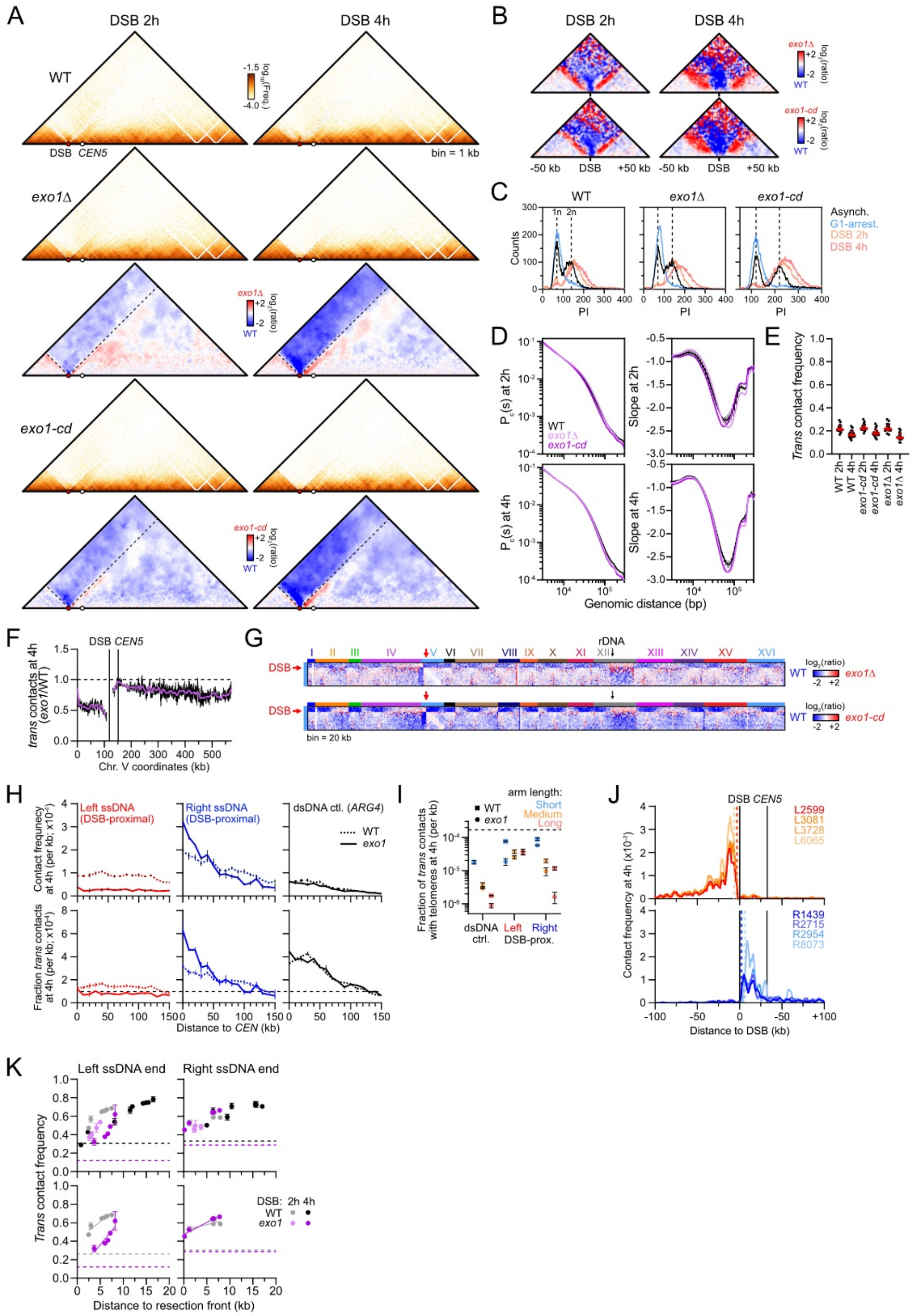


**Figure S5: Exo1 catalytic activity promotes genome-wide homology search. (Related to Figure 4)**

(A) Hi-C contact maps and Serpentine ratio maps of chr. V in WT (APY266, n = 4), *exo1Δ* (APY1160, n = 1) and *exo1-D173A* (*exo1-cd*, APY536, n = 1) cells 2 and 4 hours post-DSB induction.

(B) Ratio maps of the 100 kb region surrounding the DSB site of the data in (A). Bin: 2 kb.

(C) FACS profiles of WT (APY266), *exo1Δ* (APY1160) and *exo1-D173A* (*exo1-cd*, APY536) cells.

(D) $P_c(s)$ and derivatives of data in (A).

(E) *Trans* contact frequency determined from WT, *exo1Δ* and *exo1-cd* Hi-C data in (A). Each dot represents a chromosome. Bar: Median. Error bars: interquartile range.

(F) Ratio of *trans* contact frequency in Exo1-deficient cells over WT cells along chr. V at 4 hours post-DSB induction. Data represent mean ± SEM. Bin: 2 kb.

(G) Ratio maps of Exo1-deficient cells over WT cells of chr. V with the rest of the genome.

(H) Contact frequency of the DSB-proximal (< 8 kb from DSB) left ssDNA, right ssDNA, and of a control dsDNA site (*ARG4*) with *trans* centromere-flanking regions, from data of **Figure 4C**. Top: frequency of total contacts. Bottom: Fraction of *trans* contacts only. Data: mean ± SEM. Dotted line: genome average.

(I) Among *trans* contacts only, average contact frequency between DSB-proximal ssDNA sites and 30 kb telomeric regions as a function of chromosome arm length in Exo1-deficient cells 4 hours post-DSB induction. Data show mean ± SEM of 2 biological replicates. Dotted line indicates the genome average. WT data are from **Figure S3J**.

(J) Contact distribution in the DSB region for individual ssDNA sites on the left (top) and right (bottom) DSB ends in Exo1-deficient cells at 4 hours post-DSB induction. Dotted lines represent the position of individual viewpoints.

(K) *Trans* contact frequency for individual ssDNA sites on the left and right ssDNA ends as a function of the distance to the corresponding resection front in WT and Exo1-deficient cells at 2 and 4 hours post-DSB induction. Data: mean ± SEM. Dotted lines: baseline *trans* contact frequency of the dsDNA regions surrounding the resection front, lower for the detached left chromosomal fragment in *Exo1*-deficient cells, see (F-G).

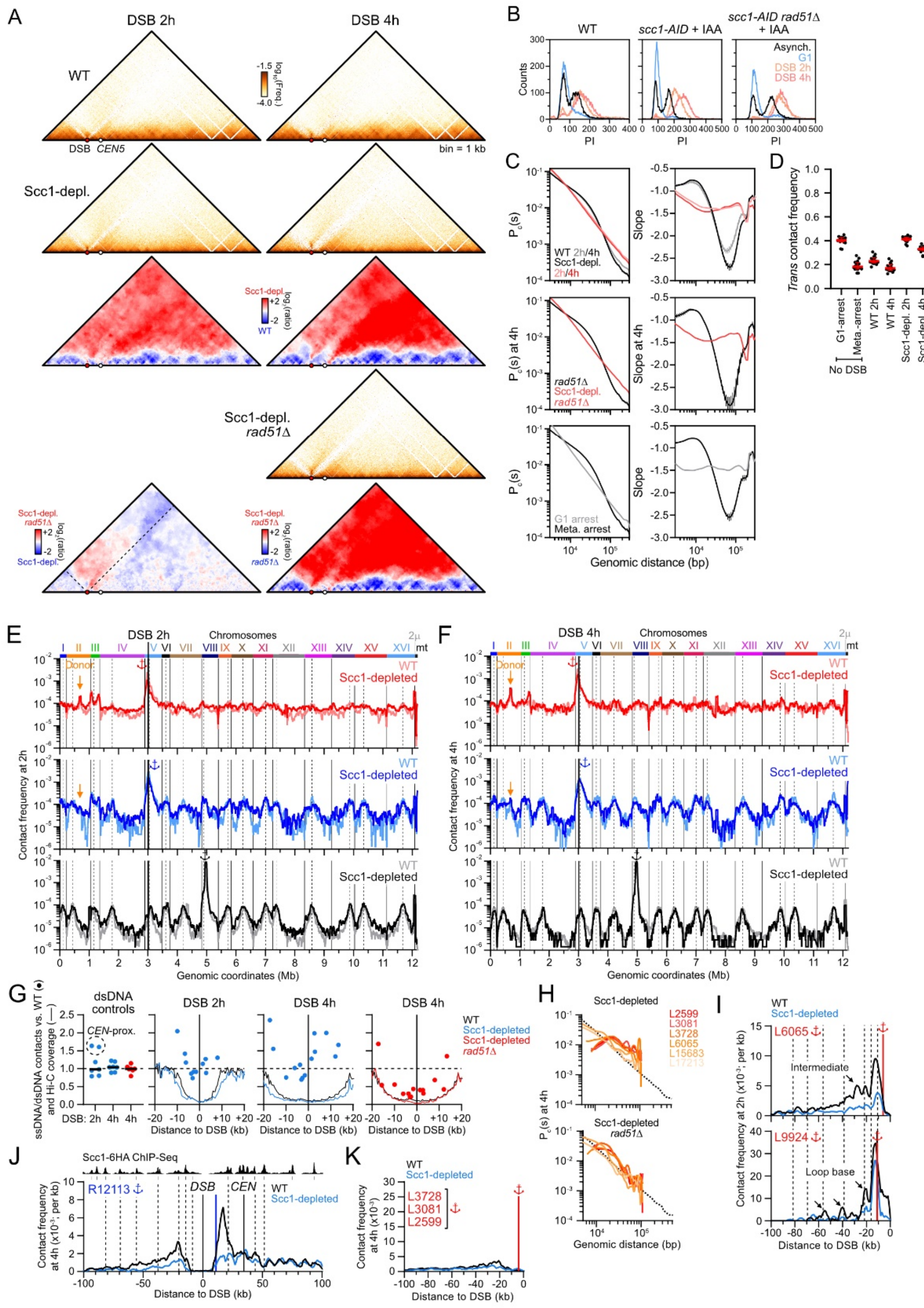

**Figure S6: Cohesin directs homology search by DSB-distal ssDNA sites in *cis*. (Related to Figure 5)**

(A) Hi-C contact maps and Serpentine ratio maps of chr. V in WT (APY266, n = 4), Scc1-depleted (APY1481, n = 1) and Scc1-depleted *rad51Δ* (APY1500, n = 1) cells at the indicated time post-DSB induction.

(B) FACS profiles of WT, Scc1-depleted (APY1481) and Scc1-depleted *rad51Δ* (APY1500) cells.

(C) $P_c(s)$ and derivatives of data in (A).

(D) *Trans* contact frequency determined from WT, Scc1-depleted and Scc1-depleted *rad51Δ* Hi-C data in (A). Each dot represents a chromosome. Bar: Median. Error bars: interquartile range.

(C-D) Undamaged G1-arrested (APY266, n = 1) and metaphase-arrested (*cdc20*-repressed; APY537, n = 2) cells are shown for comparison.

(E-F) Average genome-wide contact distribution of the left and right ssDNA ends at 2 (left) and 4 (right) hours post-DSB induction in WT (APY266) and Scc1-depleted (APY1481) cells. Other legends as in **Figure 2A**.

(G) Hi-C coverage and amount of dsDNA (left) and ssDNA (right) contacts retrieved in Scc1-depleted and Scc1-depleted *rad51Δ* cells relative to WT cells. Data: mean of individual captured ssDNA and dsDNA sites.

(H) LOWESS regression of *cis* contacts made by individual ssDNA sites on the left DSB end compared to the genome-averaged $P_c(s)$ in Scc1-depleted and Scc1-depleted *rad51Δ* cells 4 hours post-DSB induction.

(I) Side-specific *cis* contact profiles of individual ssDNA sites close to the resection front in WT and Scc1-depleted cells 2 hours post-DSB induction. Other legends as in **Figure 5D**.

(J) Contact profiles of the R12113 ssDNA site in WT and Scc1-depleted cells at 4 hours post-DSB induction on both side of the DSB region.

(K) Contact profiles of the average of 3 DSB-proximal ssDNA sites on the left DSB end in WT and Scc1-depleted cells 4 hours post-DSB induction.

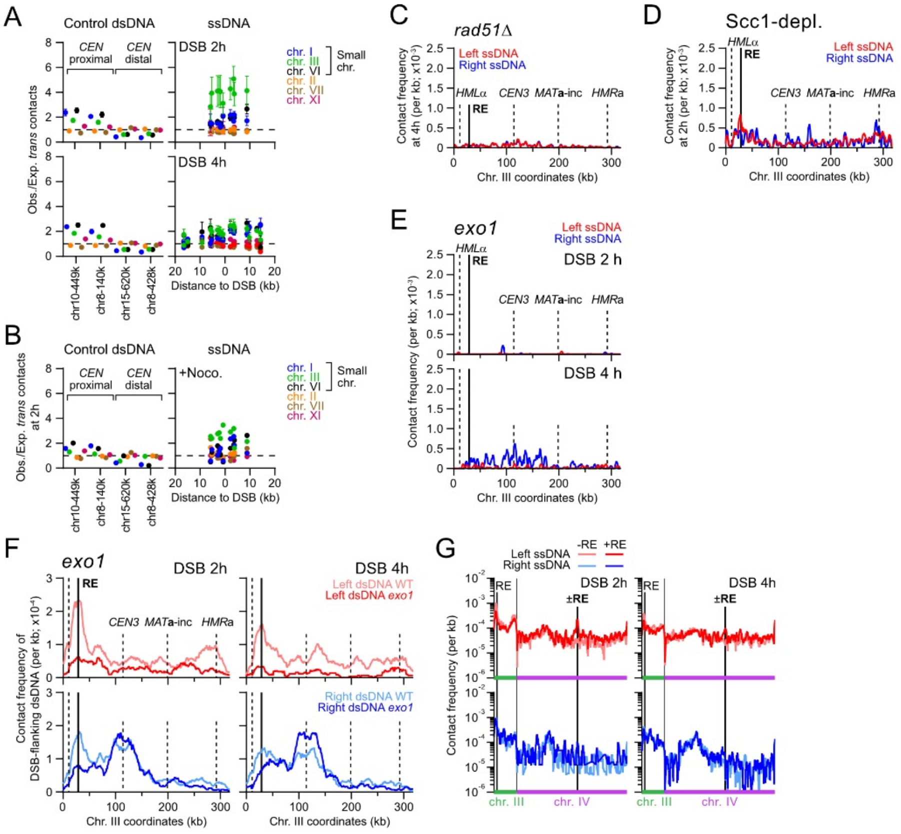


**Figure S7: The recombination enhancer stimulates early *trans* search on chr. III. (Related to Figure 6)**

(A) Enrichment of *trans* contacts per chromosome, normalized on their length, for two CEN-proximal and two CEN-distal dsDNA control sites (left) and individual ssDNA sites (right) at 2 and 4 hours post-DSB induction in WT cells. Data show mean ± SEM of 4 biological replicates. Dotted line: no enrichment. A subset of six chromosomes, including the three shortest chromosomes, is shown.

(B) Same as in nocodazole-treated WT cells 2 hours post-DSB induction (n = 1).

(C) Pondered ssHi-C contact profiles on chr. III in *rad51Δ* cells 4 hours post-DSB induction.

(D) ssHi-C contact profiles on chr. III in Scc1-depleted cells 2 hours post-DSB induction.

(E) DSB-proximal (< 4 kb from the DSB) ssHi-C contact profiles on chr. III in Exo1-deficient cells 2 and 4 hours post-DSB induction.

(F) Hi-C contact profiles of DSB-flanking dsDNA regions 5 kb away to the left and right of the DSB site in WT and Exo1-deficient cells.

(G) ssHi-C contact profiles on chr. III and IV in WT cells (APY266) and in cells bearing an additional 700 bp RE element inserted at position 845,464 on chr. IV (APY1497) at 2 and 4 hours post-DSB induction. Signal smoothed over 10 kb.

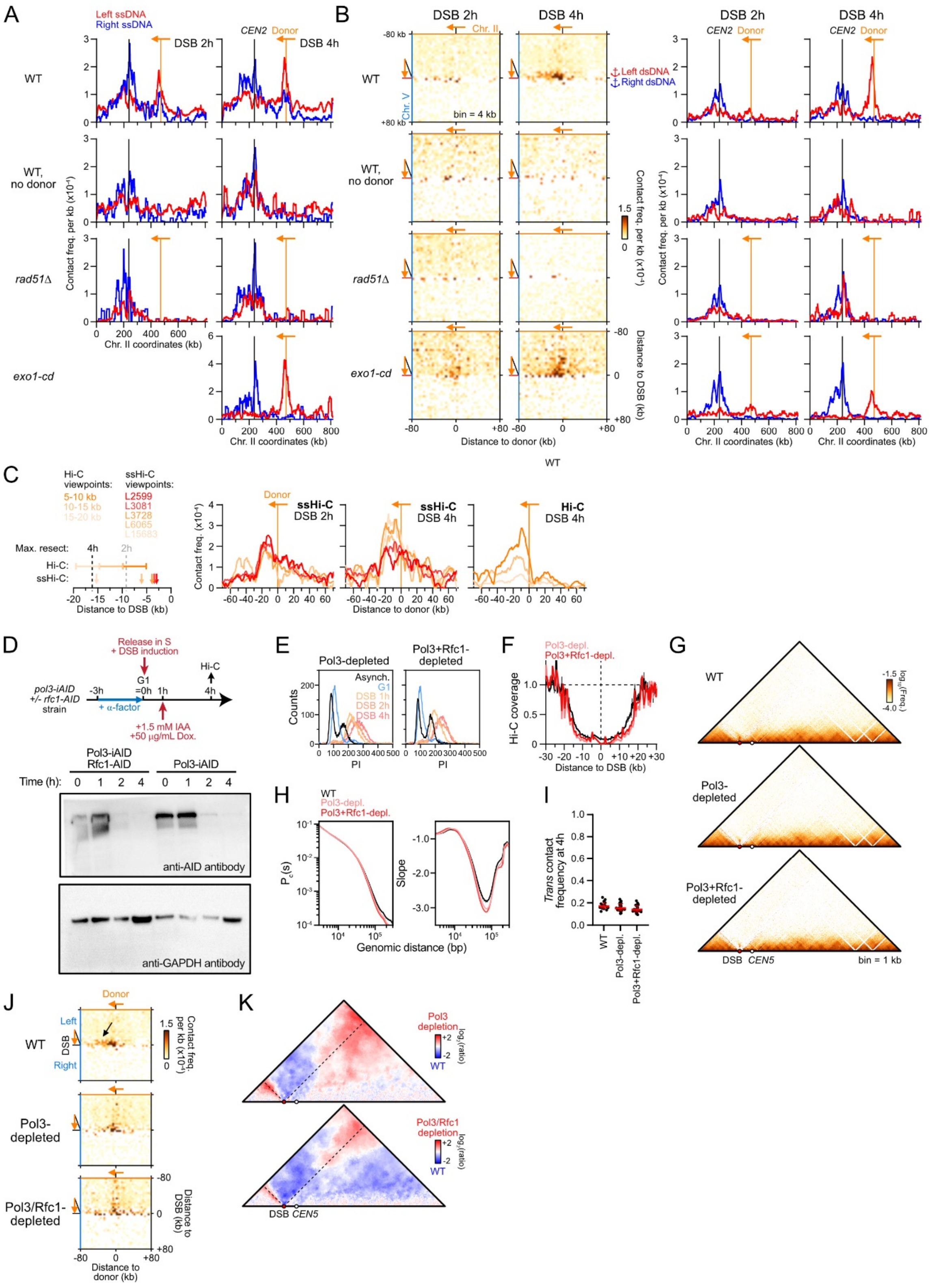


**Figure S8: Coordinated homology search by both DSB ends and side-specific NPF collapse in a donor-dependent manner. (Related to Figure 7)**

(A) ssHi-C contact profiles on chr. II in WT cells (APY266, n = 4), *rad51Δ* cells (APY679, n = 2), and exo1-D173A cells (APY536, n = 1) containing a donor at position 468 kb on chr. II and WT cells lacking a donor (APY358, n = 1) at 2 and/or 4 hours post-DSB induction.

(B) Inter-chromosomal Hi-C contact map between the DSB and the donor sites (left, bin = 4 kb) and corresponding Hi-C contact profiles of DSB-flanking dsDNA regions (right) in the same strains as in (A).

(C) Comparison of Hi-C and ssHi-C contact profiles by dsDNA regions and ssDNA sites at increasing distances from the left DSB end. The location of the viewpoints is depicted on the top left corner.

(D) Depletion procedure and Western blot analysis of Pol3-iAID alone or in combination with Rfc1-AID.

(E) FACS profiles of Pol3-depleted (APY1352) and Pol3- and Rfc1-depleted (APY1350) cells.

(F) Hi-C coverage in the DSB-surrounding region 4 hours post-DSB induction in WT, Pol3-depleted and Pol3/Rfc1-depleted cells.

(G) Hi-C contact maps of chr. V in WT (APY266, n = 4), Pol3-depleted (APY1352, n = 1) and Pol3/Rfc1-depleted (APY1350, n = 1) cells 4 hours post-DSB induction.

(H) $P_c(s)$ and its derivatives of data in (F).

(I) *Trans* contact frequency for individual chromosomes of data in (F).

(J) Inter-chromosomal Hi-C contact map between the DSB and the donor sites in WT (APY266), Pol3-depleted (APY1352) and Pol3- and Rfc1-depleted (APY1350) cells 4 hours post-DSB induction. Bin: 4 kb. Arrow: contact enrichment in the direction of D-loop extension lost in Pol3/Rfc1-depleted cells.

(K) Ratio map of chr. V in Pol3-depleted (top) and Pol3- and Rfc1-depleted (bottom) over WT cells.

**Supplementary Table 1: Genotype of the haploid *Saccharomyces cerevisiae* strains used in this study.**

| Strain | Relevant genotype | Source |
|---|---|---|
| APY266 | *MAT* **a**-inc, *ura3::LY-HOcs, lys2::LY, trp1::GAL-HO-hphMX, his3-11,15, can1-100, leu2-3,112, ade2-1, RAD5* | Piazza *et al.* 2021 |
| APY679 | *MAT* **a**-inc, *ura3::LY-HOcs, lys2::LY, trp1::GAL-HO-hphMX, his3-11,15, can1-100, leu2-3,112, ade2-1, RAD5,* ***rad51::kanMX*** | This study |
| APY1160 | *MAT* **a**-inc, *ura3::ura3::LY*-HOcs (contains mutations), lys2::LY, trp1::GAL-HO-hphMX, can1-100, ade2-1, leu2-3,112, his3-11,15, RAD5,* ***exo1::KMX*** | This study |
| APY536 | *MAT* **a**-inc, *ura3::LY-HOcs, lys2::LY, trp1::GAL-HO-hphMX, can1-100, ade2-1, leu2-3,112, his3-11,15, RAD5,* ***exo1-D173A::TRP1*** | Piazza *et al.* 2021 |
| APY1481 | *MAT* **a**-inc, *ura3::LY-HOcs, lys2::LY, trp1::GAL-HO-hphMX, his3::pADH1-OsTIR1-9Myc::HIS3, can1-100, leu2-3,112, ade2-1, RAD5,* ***SCC1-PK3-AID::kanMX*** | This study |
| APY1500 | *MAT* **a**-inc, *ura3::LY-HOcs, lys2::LY, trp1::GAL-HO-hphMX, his3::pADH1-OsTIR1-9Myc::HIS3, can1-100, leu2-3,112, ade2-1, RAD5,* ***SCC1-PK3-AID::kanMX, rad51::kanMX*** | This study |
| APY513 | *MAT* **a**-inc, *ura3::LY-HOcs, lys2::LY, trp1::GAL-HO-hphMX, can1-100, his3::pADH1-OsTIR1-9Myc::HIS3, ade2-1, leu2-3,112, RAD5,* ***CDC45-FlagX5-mini-AID::kanMX*** | Piazza *et al.* 2021 |
| APY1548 | *MAT* **a***-inc, ura3::LY-HOcs, lys2::LY, trp1::GAL-HO-hphMX, his3-11,15, leu2-3,112, can1-100, ade2-1, RAD5,* ***RE Δ*** | This study |
| APY1497 | *ura3::LY-HOcs, lys2::LY, can1-100, trp1::GAL-HO-hphMX, ade2-1 his3-11,15 leu2-3,112 RAD5,* ***chrIV-845464::RE(700 bp)*** | This study |
| APY358 | *MAT* **a**-inc, *ura3::LY-HOcs,* ***lys2::URA3,*** *trp1::GAL-HO-hphMX, his3-11,15, can1-100, leu2-3,112, ade2-1, RAD5* | This study |
| APY1350 | *MAT* **a**-inc, *ura3::LY-HOcs, lys2::LY, trp1::GAL-HO-hphMX, his3-11,15, can1-100, leu2-3,112, ade2-1, RAD5,* ***pol3-iAID (KanMX N-term & hphMX C-term), RFC1-AID-9Myc-hphMX,*** *SSN6::pST1760 (TetR, OsTir1)::HIS3* | This study |
| APY1352 | *MAT* **a**-inc, *ura3::LY-HOcs, lys2::LY, trp1::GAL-HO-hphMX, his3-11,15, can1-100, leu2-3,112, ade2-1, RAD5,* ***pol3-iAID (KanMX N-term & hphMX C-term),*** *SSN6::pST1760 (TetR, OsTir1)::HIS3* | This study |

**Supplementary Table 2**: **DNA oligonucleotides used in this study.**

Tab 1: Annealing oligonucleotides:

| **Name** | **Orientation** | **DpnII_to_HOcs _distance** | **S288c_DSB_ LY_ genome DpnII_site_st art** | **Annealing_oligonucleotide_sequence** |
|---|---|---|---|---|
| Probe_URA-L-17213-MfeI-RC_v5 | - | -17213 | 101550 | ACcATACAtTTGTCTACTTTAACAATAGAAGTTtATAAGGTTGGGCCTCTTTTAAGAGaACAGGTATAGT**GATC**TCATCG |
| Probe_URA-L-16220-MfeI-RC_v5 | - | -16220 | 102543 | CCATGA**GATC**TTGCACCTTTTcGTTATGCCTAAATTACTACAGCTaAATTGAcAATACATGTTTGTGCATTTACAGATGC |
| Probe_URA-L-15683-SspI-RC | - | -15683 | 103080 | GCGAGTGATAAGAACTtTCTATTAGGTCGATTAaCAGACATTTCTTGTGCAACGTTGAAtTATTCCAATA**GATC**TTATGT |
| Probe_URA-L-9924-MfeI-RC_v5 | - | -9924 | 108839 | AAGTTT**GATC**GTAGGTATAGACtGGACAATATGCCGGAATATGTAAGGCAATTaTTCCAAGtTTTGGAAGGTATTtATTT |
| Probe_URA-L-6532-MfeI-RC | - | -6532 | 112231 | CGATGG**GATC**TTCTTGACTTgCTTCTTCTTTGGATaCTACATTTGTGCCAtTTGTAaCGGCCTTCAATACAaCATTTGTG |
| Probe_URA-L-6065-SspI-RC | - | -6065 | 112698 | TTTCAtAACTCGAGTTATTTGCGGTACCAaCATgTAAAAGAATAGAAAtTATTCTTGTTAGAGTGtGCTC**GATC**TGTAAT |
| Probe_URA-L-4543-SspI-RC | - | -4543 | 114220 | ATCAAGTCTTCGTACAATGTTAGTATAtTATTAACAATGTTTAACCATAATTCCTTGGAGAAAATTTTAG**GATC**TTTGTT |
| Probe_URA-L-3728-SspI-RC | - | -3728 | 115035 | ATGAAATTTCtTCTAATAGTCCTAGGACaCACATGAAGTaCTCATTTGTCAAAtTATTGCTGTCTGTCTT**GATC**GATTaA |
| Probe_URA-L-3081-MfeI-RC | - | -3081 | 115682 | TGTATTGTCATaAATTGGCGCAGTAGCCTCAATTTCAACGTCGTTTGcCTCTGGTGTTTGTTaATGTGCA**GATC**CATGAG |
| Probe_URA-L-2599-MfeI-RC | - | -2599 | 116186 | GTTAAATCCAAAACGGCAGAAGCCTGAATGAAACATATtAACCAtTTGGAGtACGTCAATaAATTCTGGG**GATC**CATTGC |
| Probe_URA-R-1439-SspI | + | 1439 | 120223 | TTGGCA**GATC**TTTAACTCGGCTTTAGTTATaCAAGTTACTTGCAtTATTTCCTTCTGCGAGAGTACATTTGCCCTAAAC |
| Probe_URA-R-2715-93-SspI_v5 | + | 2715 | 121510 | CCAATAACAtTCAGAATAcTTTTATTTTTATGTTGCAAAATAAGCTcTACGACTTTTTT**GATC**TTCGACG**GATC**AGGGCC |
| Probe_URA-R-2954-SspI_v5 | + | 2954 | 121738 | GTAtTCATGTCAtTATTGTCAGGGGTTAACTTTCCGGTAAACTTCAAAAATaGGGACAAAAGACTAACTT**GATC**CAACTC |
| Probe_URA-R-8073-SspI | + | 8073 | 126857 | TATCGTCATAcCTGTGCTTTCTGTTAcCGTATTGGAAtTATTTCCAGCTaGtGTCGAAAaCTTCTTTCAG**GATC**ACGTGA |
| Probe_URA-R-9028-MfeI_v5 | + | 9028 | 127812 | GTATTT**GATC**TATTCGCAtTTGTCCGGGGCaCATTATgAATAGGTTTTGGGCACTTCGAATATAAAAGCAACTCAATGAG |
| Probe_URA-R-10241-SspI_v5 | + | 10241 | 129025 | ATC**G**AT**GATC**CCAAATACCTACCtATAACTCAGTCTTTGAtTATTACACATTTAATTCAgTCTGCAACTAACTCTCaATC |
| Probe_URA-R-12116-SspI | + | 12116 | 130900 | TACTGAAAAATACGTCCGTCAaGTCTCTAGAGAGGTACTGGAAaCCcTCTTAATAcTATTACAAAGCCAA**GATC**CACAGA |
| Probe_URA-R-13482-SspI_v5 | + | 13482 | 132266 | ACACAG**GATC**TGAACATaAACCTGTAGAGaATGCAAGTTTGGAATTGTATAtTATTACTCAACAtATTTTACAATTTTTA |
| Probe_URA-R-14040-MfeI | + | 14040 | 132824 | CGTTTTTAGAATATATTGTAATAAAACACAATTaATAATACAGTTgTCTCTTCGTCCTCTtATGGTGGGA**GATC**TATTAA |
| Probe_URA-L-774-MfeI-RC | - | -774 | 118011 | ATATCATCTTGAGTAGGGACATACAtTTGGGCACCTAAAAATAATGGTGTAAACATtTCTCTTTaAATTG**GATC**ATGTGC |
| Probe_URA-L-1560-SspI-RC | - | -1560 | 117225 | TGGATG**GATC**GCTTAGCGCAGCAGTCAAATAgTGAGTAAATTGGTgCGCAACAATGGTTACTCTTTCATTCGAATACAGT |

Nucleotide in lower case mark SNPs relative to the reference sequence. DpnII sites are in bold.

Tab 2: Capture oligonucleotides

| Name | Orientation | Chromosome | Start_S288c_DSB_LY_Capture | End_S288c_DSB_LY_Capture | Capture oligo 5'-biot (60 mers) |
|---|---|---|---|---|---|
| Probe_URA-L-17213-MfeI-RC | - | chr5 | 101554 | 101614 | TGTCTACTTTAACAATAGAAGTTtATAAGGTTGGGCCTCTTTTAAGAGaACAGGTATAGT |
| Probe_URA-L-16220-MfeI-RC | - | chr5 | 102483 | 102543 | TTGCACCTTTTcGTTATGCCTAAATTACTACAGCTaAATTGAcAATACATGTTTGTGCAT |
| Probe_URA-L-15683-SspI-RC | - | chr5 | 103084 | 103144 | AGAACTtTCTATTAGGTCGATTAaCAGACATTTCTTGTGCAACGTTGAAtTATTCCAATA |
| Probe_URA-L-9924-MfeI-RC | - | chr5 | 108779 | 108839 | GTAGGTATAGACtGGACAATATGCCGGAATATGTAAGGCAATTaTTCCAAGtTTTGGAAG |
| Probe_URA-L-6532-MfeI-RC | - | chr5 | 112171 | 112231 | TTCTTGACTTgCTTCTTCTTTGGATaCTACATTTGTGCCAtTTGTAaCGGCCTTCAATAC |
| Probe_URA-L-6065-SspI-RC | - | chr5 | 112712 | 112772 | TTTCAtAACTCGAGTTATTTGCGGTACCAaCATgTAAAAGAATAGAAAtTATTCTTGTTA |
| Probe_URA-L-3728-SspI-RC | - | chr5 | 115039 | 115099 | tTCTAATAGTCCTAGGACaCACATGAAGTaCTCATTTGTCAAAtTATTGCTGTCTGTCTT |
| Probe_URA-L-3081-MfeI-RC | - | chr5 | 115686 | 115746 | TaAATTGGCGCAGTAGCCTCAATTTCAACGTCGTTTGcCTCTGGTGTTTGTTaATGTGCA |
| Probe_URA-L-2599-MfeI-RC | - | chr5 | 116200 | 116260 | ATTGACGTaCTCCAAaTGGTTaATATGTTTCATTCAGGCTTCTGCCGTTTTGGATTTAAC |
| Probe_URA-R-1439-SspI | + | chr5 | 120227 | 120287 | TTTAACTCGGCTTTAGTTATaCAAGTTACTTGCAtTATTTCCTTCTGCGAGAGTACATTT |
| Probe_URA-R-2715-93-SspI | + | chr5 | 121440 | 121500 | CCAATAACAtTCAGAATAcTTTTATTTTTATGTTGCAAAATAAGCTcTACGACTTTTTTG |
| Probe_URA-R-2954-SspI | + | chr5 | 121678 | 121738 | CAtTATTGTCAGGGGTTAACTTTCCGGTAAACTTCAAAAATGGGGACAAAAGACTAACTT |
| Probe_URA-R-8073-SspI | + | chr5 | 126797 | 126857 | cCTGTGCTTTCTGTTAcCGTATTGGAAtTATTTCCAGCTaGtGTCGAAAaCTTCTTTCAG |
| Probe_URA-R-9028-MfeI | + | chr5 | 127816 | 127876 | TATTCGCAtTTGTCCGGGGCaCATTATgAATAGGTTTTGGGCACTTCGAATATAAAAGCA |
| Probe_URA-R-10241-SspI | + | chr5 | 129029 | 129089 | CCAAATACCTACCtATAACTCAGTCTTTGAtTATTACACATTTAATTCAgTCTGCAACTA |
| Probe_URA-R-12116-SspI | + | chr5 | 130840 | 130900 | TACGTCCGTCAaGTCTCTAGAGAGGTACTGGAAaCCcTCTTAATAcTATTACAAAGCCAA |
| Probe_URA-R-13482-SspI | + | chr5 | 132270 | 132330 | TGAACATaAACCTGTAGAGaATGCAAGTTTGGAATTGTATAtTATTACTCAACAtATTTT |
| Probe_URA-R-14040-MfeI | + | chr5 | 132754 | 132814 | ATATATTGTAATAAAACACAATTaATAATACAGTTgTCTCTTCGTCCTCTtATGGTGGGA |
| Probe_URA-L-1560-SspI-RC | - | chr5 | 117165 | 117225 | GCTTAGCGCAGCAGTCAAATAgTGAGTAAATTGGTgCGCAACAATGGTTACTCTTTCATT |
| chr4-64420-CDC13 | NA (dsDNA) | chr4 | 64420 | 64480 | CAACTCACTTGTGGATATCTTCAACAATTTAATAGAAATGAATAGAGACGAGAAAAACAG |
| chr8-428677-DNA2 | NA (dsDNA) | chr8 | 428677 | 428737 | CTGATGACTACGAAGATGTCCAAAATCCCTCTTCTACACCTATAGTCCCTAATCGACTGA |
| chr15-996452-MEK1 | NA (dsDNA) | chr15 | 996452 | 996512 | ATGAGACCGTTGTATAGCTGCAACCTTGCAACCAAAGATGATATTGAGATGGCAGGTGGC |
| chr2-650593-MET8 | NA (dsDNA) | chr2 | 650593 | 650653 | CCGAACTGGGACCCCACGAAGAATGAAATTTACGAGTACATCCGCAGTGACTTCAAAGAC |
| chr15-620339-PDR5 | NA (dsDNA) | chr15 | 620339 | 620399 | CTAAAGAAACCAATACTTTCCAAATCTTGAAACCAATGGATGGTTGCCTAAACCCAGGTG |
| chr10-449622-POL31 | NA (dsDNA) | chr10 | 449622 | 449682 | GGTGGTTCTGATGAAATTATGCTCGAAGATGAAAGTGGGAGAGTGCTTCTAGTGGGAGAT |
| chr8-140404-ARG4 | NA (dsDNA) | chr8 | 140404 | 140459 | AAATAATGGCTCTTTGTCTTCTTGCATGTCTTTATCATAAGTAGATGGGATACCCTTCAA |

#### Tab 3: Genome editing

| RE deletion (chrIII:29107-30801) | Sequence (5'-3') |
|---|---|
| Cas9 guide left | ACAACGGAATCATTGAGCGT |
| Cas9 guide right | CAACGGAATCATTGAGCGA |
| Repair template | AGCATCAACACATAAATCCTTGCTTAGCTCAATTAAATATACTAGTAAATAAGTATATAAACAATAATTTTGCATTTTATTTTACTGGAAACTCTTCTCAAAACCAAATTGCGCAAGGATTGATTCAGTACAATTATGCAAACTCGAAAAGTAAATAAACAAAAAGATACA/CTTATAAAATGAATATTTCAATTGATGAATAGCTATATATTGGATACAAAAATTAGCATTTAATCGAAAACTGCAGCATGTATTTAATCGAAAACTACAGCATGTAGCTATGATACGACAGAAGATTTTTGTTTTTATAGTTAAGTCAAGAAGAAATTCTATTTGTCCAGCAATCCGGCGCAAAGAAGACTACTAAAGGG |

| RE insertion (chrIV:845464) | Sequence (5'-3') |
|---|---|
| RE-atchrIV-845464-F | TAACTAGGATTTAATCATAAAAAAAAAAGTGATTGACACTGTTATTCAGTTCTGGACTATCGTATAGAGTTGAGTGAAAGGTAAATAAAC |
| RE-atchrIV-845464-R | TGGGCTTGGTAATCGTTGAAAAATTTTCACATATTTTATGAACAGCTGAGCCTATTAACAATGAAAATAGACCGAAGTACAACAATATCC |
